# The Alzheimer’s Disease Metabolome: Effects of Sex and *APOE* ε4 genotype

**DOI:** 10.1101/585455

**Authors:** Matthias Arnold, Kwangsik Nho, Alexandra Kueider-Paisley, Tyler Massaro, Barbara Brauner, Siamak MahmoudianDehkordi, Gregory Louie, M. Arthur Moseley, J. Will Thompson, Lisa St John Williams, Jessica D. Tenenbaum, Colette Blach, Rui Chang, Roberta D. Brinton, Rebecca Baillie, Xianlin Han, John Q. Trojanowski, Leslie M. Shaw, Michael W. Weiner, Eugenia Trushina, Jon B. Toledo, Jan Krumsiek, P. Murali Doraiswamy, Andrew J. Saykin, Rima Kaddurah-Daouk, Gabi Kastenmüller, for the Alzheimer’s Disease Neuroimaging Initiative, the Alzheimer’s Disease Metabolomics Consortium

**Affiliations:** Department of Psychiatry and Behavioral Sciences, Duke University, Durham, NC, USA; Institute of Bioinformatics and Systems Biology, Helmholtz Zentrum München, German Research Center for Environmental Health, Neuherberg, Germany; Department of Radiology and Imaging Sciences and the Indiana Alzheimer Disease Center, Indiana University School of Medicine, Indianapolis, IN, USA; Duke Clinical Research Institute, Duke University, Durham, NC, USA; Duke Proteomics and Metabolomics Shared Resource, Center for Genomic and Computational Biology, Duke University, Durham, NC, USA; Department of Biostatistics and Bioinformatics, Duke University, Durham, NC, USA; Duke Molecular Physiology Institute, Duke University, Durham, NC, USA; Center for Innovation in Brain Science, University of Arizona, Tucson, AZ, USA; Departments of Pharmacology and Neurology, College of Medicine, University of Arizona, Tucson, AZ, USA; Rosa & Co LLC, San Carlos, CA, USA; University of Texas Health Science Center at San Antonio, San Antonio, TX, USA; Department of Pathology & Laboratory Medicine, University of Pennsylvania, Philadelphia, PA, USA; Center for Imaging of Neurodegenerative Diseases, Department of Radiology, San Francisco VA Medical Center/University of California San Francisco, San Francisco, CA, USA; Departments of Neurology and Molecular Pharmacology and Experimental Therapeutics, Mayo Clinic, Rochester, MN, USA; Department of Neurology, Houston Methodist Hospital, Houston, TX, USA; Institute for Computational Biomedicine, Englander Institute for Precision Medicine, Department of Physiology and Biophysics, Weill Cornell Medicine, New York, NY, USA; Duke Institute of Brain Sciences, Duke University, Durham, NC, USA; Department of Medicine, Duke University, Durham, NC, USA; German Center for Diabetes Research (DZD), Neuherberg, Germany

## Abstract

Recent studies have provided evidence that late-onset Alzheimer’s disease (AD) can in part be considered a metabolic disease. Besides age, female sex and *APOE* ε4 genotype represent strong risk factors for AD. They also both give rise to large metabolic differences, suggesting that metabolic aspects of AD pathogenesis may differ between males and females and between *APOE* ε4 carriers and non-carriers. We systematically investigated group-specific metabolic alterations by conducting stratified association analyses of 140 metabolites measured in serum samples of 1,517 AD neuroimaging initiative subjects, with AD biomarkers for Aβ and tau pathology and neurodegeneration. We observed substantial sex differences in effects of 15 metabolites on AD biomarkers with partially overlapping differences for *APOE* ε4 status groups. These metabolites highlighted several group-specific alterations not observed in unstratified analyses using sex and *APOE* ε4 as covariates. Combined stratification by both variables uncovered further subgroup-specific metabolic effects limited to the group with presumably the highest AD risk: *APOE* ε4+ females. Pathways linked to the observed metabolic alterations suggest that females experience more expressed impairment of mitochondrial energy production in AD than males. These findings indicate that dissecting metabolic heterogeneity in AD pathogenesis may enable grading of the biomedical relevance of specific pathways for specific subgroups. Extending our approach beyond simple one- or two-fold stratification may thus guide the way to personalized medicine.

**Significance statement:** Research provides substantial evidence that late-onset Alzheimer’s disease (AD) is a metabolic disease. Besides age, female sex and APOEε4 genotype represent strong risk factors for AD, and at the same time give rise to large metabolic differences. Our systematic investigation of sex and APOE ε4 genotype differences in the link between metabolism and measures of pre-symptomatic AD using stratified analysis revealed several group-specific metabolic alterations that were not observed without sex and genotype stratification of the same cohort. Pathways linked to the observed metabolic alterations suggest females are more affected by impairment of mitochondrial energy production in AD than males, highlighting the importance of tailored treatment approaches towards a precision medicine approach.

## 1. Introduction

Female sex is regarded a major risk factor for Alzheimer’s disease (AD). Of 5.3 million people in the United States diagnosed with AD at age 65+, more than 60% are women. The lifetime risk of developing AD at age 45 may be almost double in females than in males (1), though the role and magnitude of sexual dimorphism in predisposition and progression to AD are controversial (2). While age is the strongest risk factor for late-onset AD (LOAD), women’s higher life expectancy only partially explains the observed sex difference in frequency and lifetime risk. Complexity is added by studies showing a significant sex difference in effects of the *APOE* ε4 genotype, the strongest common genetic risk factor for LOAD. These studies report higher risk estimates for female ε4 carriers, a finding that seems to be also age dependent (3, 4). *APOE* ε4 is also associated with AD biomarkers in a sex-dependent way with larger risk estimates for women (3, 5, 6), although these findings are not consistent (5). Also, studies suggest that sex differences in AD may change during the course of disease (7). Overall risk for mild cognitive impairment (MCI), the prodromal stage of AD is higher in males (8), while progression to AD occurs faster in females, at least partly in *APOE* ε4-dependent ways (7, 9). The mechanisms underlying this sex-linked and partly intertwined *APOE* ε4- and age-dependent heterogeneity in AD susceptibility and severity are only beginning to unravel, calling for novel approaches to further elucidate molecular sex differences in AD risk and biomarker profiles.

All of the aforementioned major AD risk factors have profound effects on metabolism (10–13), supporting the view of AD as a metabolic disease. Recent availability of high-throughput metabolomics techniques, which can measure hundreds of small biochemical molecules (metabolites) simultaneously, enabled the study of metabolic imprints of age, genetic variation, and sex very broadly, covering the entire metabolism: (a) Age-dependent differences were observed in phosphatidylcholines (PCs), sphingomyelins (SMs), acylcarnitines, ceramides, and amino acids (12). (b) As expected from APOE’s role in cholesterol and lipid metabolism (14), common genetic variants in APOE were associated with blood cholesterol levels in genome- and metabolome-wide association studies (14, 15).(c) Analogous to age, sex also affects blood metabolite levels from a broad range of biochemical pathways. In a healthy elderly population females showed higher levels of most lipids except lyso-PCs. Levels of most amino acids—including branched chain amino acids (BCAAs)—were higher in males, though glycine and serine levels were higher in women (10, 11). In women, higher levels of ether-PC and SM species were associated with longevity; no significant differences were observed in men. Thus, based on results from large-scale metabolomics studies, aging may influence a wider range of metabolites in women than men, highlighting the need for sex-stratified analyses(12).

Many metabolites affected by female sex, age, and *APOE* genotype(e.g., BCAAs, glutamate, lipids) appear altered in AD independent of these risk factors (16, 17).MCI patients have alterations in lipid and lysine metabolism and the tricarboxylic acid cycle (18, 19). We identified metabolic alterations in various stages across the trajectory of AD. Higher levels of SMs and PCs were noted in early stages of AD (e.g. abnormal CSF Aβ_1-42_ levels), whereas intermediate changes (CSF total tau), were correlated with increased levels SMs levels and long-chain acylcarnitines (20). Changes in brain volume and cognition, usually noted in later stages, were correlated with a shift in energy substrate utilization from fatty acids to amino acids, especially BCAAs. Other studies reported metabolic alterations in AD which support these findings, including alterations in PCs in AD (19) and sphingolipid transport and fatty acid metabolism in MCI/AD compared to cognitively normal (CN) (21). Metabolomics analysis of brain and blood tissue revealed that bile acids, important regulators of lipid metabolism and products of human-gut microbiome co-metabolism, were altered in AD (22) and associated with CSF Aβ_1-42_, p-tau, and brain glucose metabolism and atrophy (23). In most of these studies, sex and *APOE* ε4 genotype were used as covariates. Thus, these studies may have missed sex-specific associations between AD and metabolite levels or associations with opposite effect directions for the sexes. Similarly, sex-by-*APOE* genotype interactions would be masked.

We examined the role of sex in the relationship between metabolic alterations and AD, to elucidate possible metabolic underpinnings for the observed sexual dimorphism in AD susceptibility and severity. Using metabolomics data from 1,517 subjects of the AD Neuroimaging Initiative (ADNI) cohorts, we investigated how sex modifies the associations of representative A-T-N biomarkers (24) (A: CSF Aβ_1-42_ pathology; T: CSF p-tau; N: region of interest (ROI)-based glucose uptake measured by FDG-PET) with 140 blood metabolites using stratified analyses and systematic comparison of effects between the sexes. In downstream analyses, we inspected sex-differences in metabolic effects on AD biomarkers for dependencies on *APOE* genotype, both by interaction analysis and sub-stratification.

## 2. Methods

### 2.1. Study subjects

Study data were obtained from the ADNI database (http://adni.loni.usc.edu/). For up-to-date information, see www.adni-info.org. For information on data availability and accessibility, see **Supplementary Text 1.**We included 1,517 baseline serum samples of fasting subjects pooled from ADNI-1, GO, and 2. Demographics and distributions of key risk factors are in **Table 1**. AD dementia diagnosis was established based on the NINDS-ADRDA criteria for probable AD. MCI participants did not meet these AD criteria and had largely intact functional performance, meeting predetermined criteria for amnestic MCI (25). The AMP-AD Knowledge Portal is the distribution site for data, analysis results, analytical methodology and research tools generated by the AMP-AD Target Discovery and Preclinical Validation Consortium and multiple Consortia and research programs supported by the National Institute on Aging.

**Table 1:**
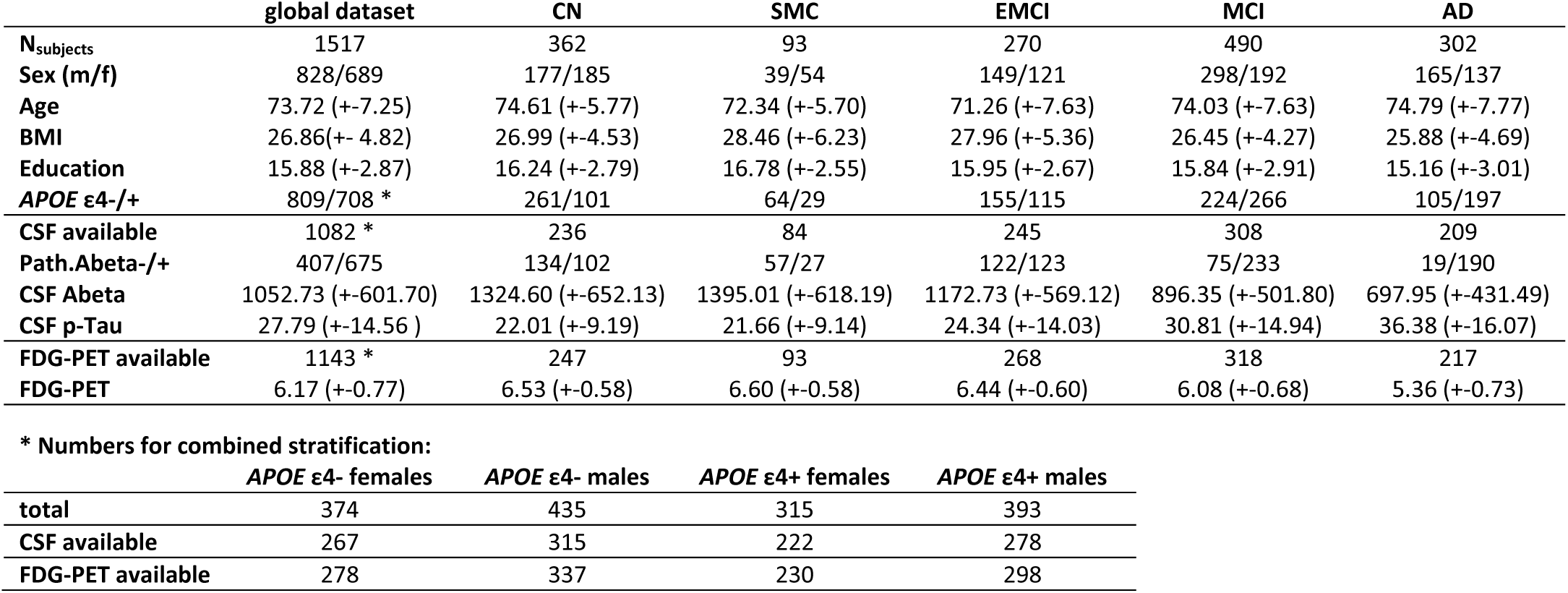
Characteristics of the 1,517 ADNI samples included in this study. CN: cognitively normal. SMC: subjective memory complaints; EMCI: early mild cognitive impairment; MCI: mild cognitive impairment; AD: probably Alzheimer’s disease; BMI: body-mass-index; *APOE* ε4-/+: non-carriers and carriers of the *APOE* ε4 allele, Path.Abeta-/+: participants who have normal and pathological CSF Abeta levels; respectively.

Data from the Rush Alzheimer’s Center (RADC) Religious Orders Study and Memory and Aging Project (ROS/MAP) (https://www.radc.rush.edu/home.htm) were used to validate key findings from ADNI. A variety of clinical, demographic, and other variables are available through RADC (https://www.radc.rush.edu/docs/var/variables.htm), while omics datasets generated from biospecimens donated by ROS and MAP participants are made available through the AMP-AD Knowledge Portal.

### 2.2. Metabolomics data acquisition

Metabolites were measured with the targeted AbsoluteIDQ-p180 metabolomics kit (BIOCRATES Life Science AG, Innsbruck, Austria), with an ultra-performance liquid chromatography (UPLC)/MS/MS system [Acquity UPLC (Waters), TQ-S triple quadrupole MS/MS (Waters)] which provides measurements of up to 186 endogenous metabolites. Sample extraction, metabolite measurement, identification, quantification, and primary quality control (QC) followed standard procedures (20, 26).

### 2.3. Metabolomics data processing

Metabolomics data processing followed the protocol previously described (20, 26) with a few adjustments. In brief, raw metabolomics data for 182 metabolites was available for 1,681 serum study samples and, for each plate, 2-3 NIST Standard Reference samples were available. Furthermore, we also had blinded duplicated measurements for 19 samples (ADNI-1) and blinded triplicated measurements for 17 samples (ADNI-GO and −2) distributed across plates. We first excluded 22 metabolites with large numbers of missing values (> 40%). Then, we removed plate batch effects using cross-plate mean normalization using NIST metabolite concentrations. Duplicated and triplicated study samples were then used to calculate the coefficients of variation (exclusion criterion >20%) and intra-class correlation (exclusion criterion <0.65) for each metabolite. We removed 20 metabolites that violated these thresholds. Next, we excluded non-fasting samples (n=108), imputed missing metabolite data using half the value of the lower limit of detection per metabolite and plate, log2-transformed metabolite concentrations, centered and scaled distributions to a mean of zero and unit variance and winsorized single outlying values to 3 standard deviations. We then used the Mahalanobis distance for detection of multivariate subject outliers, applying the critical Chi-square value for *P* < 0.01 and removing 42 subjects. Finally, metabolites were adjusted for significant medication effects using stepwise backwards selection (for details see (26)). The final QC-ed metabolomics dataset was further restricted to subjects with data on all significant covariates (Section 2.4), resulting in the study dataset of 140 metabolites and 1,517 subjects.

### 2.4. Phenotype data and covariate selection

We limited association analyses of metabolites with AD to early detectable endophenotypes, specifically to the pathological threshold for CSF Aβ_1-42_, levels of phosphorylated tau protein in the CSF (p-tau), and brain glucose metabolism measured by [^18^F] fluorodeoxyglucose (FDG)-PET. Baseline data on these biomarkers for ADNI-1, -GO, and −2 participants was downloaded from the LONI online portal at https://ida.loni.usc.edu/. For CSF biomarker data, we used the dataset generated using the validated and highly automated Roche Elecsys electrochemiluminescence immunoassays (27, 28). For FDG-PET, we used an ROI-based measure of average glucose uptake across the left and right angular, left and right temporal and bilateral posterior cingulate regions derived from preprocessed scans (co-registered, averaged, standardized image and voxel size, uniform resolution) and intensity-normalized using a pons ROI to obtain standard uptake value ratio (SUVR) means (29, 30). The pathological CSF Aβ_1-42_ cut-point (1,073 pg/ml) as reported by the ADNI biomarker core for diagnosis-independent mixture modeling (see http://adni.loni.usc.edu/methods/, accessed Oct 2017) was used for categorization since CSF Aβ_1-42_ concentrations were not normally distributed. Processed FDG-PET values were scaled and centered to zero mean and unit variance prior to association analysis; p-tau levels were additionally log2-transformed. Furthermore, we extracted covariates including age, sex, body-mass index (BMI), *APOE* ε4 genotype copies, and years of education. Covariates were separated into forced-in (age, sex, ADNI study phase, *APOE* ε4 copies) and covariates (BMI, education) selectable by backwards selection. ADNI study phase was included to adjust for remaining metabolic differences between batches (ADNI-1 and ADNI-GO/-2 were processed in separate runs) and differences in PET imaging technologies.

### 2.5. Association analyses

Association analyses of the three AD biomarkers with metabolite levels were conducted using standard linear (p-tau, FDG-PET) and logistic (pathological Aβ_1-42_) regression. For pathological CSF Aβ_1-42_, only BMI was additionally selected; for p-tau and FDG-PET the full set of covariates was used. The stratification variables sex and *APOE* ε4 copies were excluded as covariates in the sex-stratified and *APOE* ε4+/- status-stratified analyses, respectively. For identifying metabolic sex-differences, we used linear regression with metabolite levels as the dependent variable and age, sex, BMI, ADNI study phase, and diagnostic group as explanatory variables and retrieved statistics for sex. To adjust for multiple testing, we accounted for the significant correlation structure across the 140 metabolites and using Li and Ji’s method (31) determined the number of independent metabolic features (i.e. tests) to be 55, leading to a threshold of Bonferroni significance of 9.09 × 10^−4^. To assess significance of heterogeneity between strata, we used the methodology of (11, 32) that is similar to the determination of study heterogeneity in inverse-weighted meta-analysis. We further provide a scaled index of percent heterogeneity similar to the *I^2^* statistic (33).

## 3. Results

We used CSF biomarkers, FDG-PET imaging, and metabolomics data on 140 metabolites to investigate metabolic effects in relation to sex and AD and their interaction. Of 1,517 ADNI subjects, 1,082 had CSF Aβ_1-42_ and p-tau levels and 1,143 had FDG-PET data available (**Table 1**). We included all subjects with respective data regardless of diagnostic classification, as we were interested in these three representatives of the A-T-N AD biomarker schema (24) as our main readouts. In this data set, there was no significant difference in the number of *APOE* ε4+/- subjects between the sexes (*P* > 0.3). Of the three AD biomarkers, only p-tau levels were significantly different between sexes (corrected *P* = 0.01) with slightly higher levels in females.

Previous studies consistently showed widespread metabolic sex-differences, metabolic imprint of genetic variance in the *APOE* locus, and significant associations between blood metabolites and AD biomarkers independent of (i.e. adjusted for) sex. The current study added specific examination of the following central questions (**Supplementary Figure 1**): (a) Are metabolic sex-differences changed due to presence of (probable) AD?, (b) Are metabolite associations with A-T-N biomarkers modified by sex?, and (c) Is there evidence for *APOE* ε4 status influencing metabolite associations with A-T-N biomarkers that show differences between sexes?

### 3.1. Metabolic sex-differences are unaffected by MCI and probable AD status

We tested whether sex-associated differences in blood metabolite levels differ between patients with probable AD, subjects with late MCI, and CN subjects in ADNI. In the complete cohort (n = 1,517), 108 of 140 metabolites were significantly associated with sex after multiple testing correction while adjusting for age, BMI, ADNI study phase, and diagnostic group. Seventy of these associations replicate previous findings in a healthy population using a prior version of the same metabolomics platform (11) that provides measurements on 92 of the 108 metabolites identified in ADNI. All SMs and the majority of PCs were more abundant in women. The majority of biogenic amines, amino acids and acylcarnitines were more abundant in men.

Stratifying subjects by diagnostic group revealed that 53 of the 108 metabolites showing significant sex-differences were also significant in each of the three groups (AD, MCI, CN) alone, while 14 showed no significant difference in any group, probably due to lower statistical power after stratification (Supplementary Table 1, **Supplementary Figure 2**). Significant sex-differences limited to one diagnostic group were found for 8 metabolites (PC aa C34:1, PC ae C34:3, PC ae C36:3, PC ae C36:4, PC ae C38:5, PC ae C40:5, Histidine, C6/C4:1-DC) in patients with probable AD, for 7 metabolites (C0, C3, C9, C18:2, SDMA, Spermidine, t4-OH-Pro) in MCI, and for 6 metabolites (PC aa C42:0, PC ae C32:1, PC ae C42:3, SM(OH) C24:1, Sarcosine, Aspartate) in CN, although all significant sex-differences found were also significant in the full cohort. Comparisons of beta estimates for sex between AD and CN showed no significant effect heterogeneity, indicating reduced power as source for these observed differences. Only PC aa C34:1 showed significant (*P* = 0.029) heterogeneity between AD compared to CN. Interestingly, in the larger healthy reference cohort, sex did not significantly affect the blood metabolite levels when adjusting for the study covariates (i.e., age and BMI) (11). In summary, we found that sex differences of blood metabolite levels are consistent (if we neglect the reduced power due to stratification) across diagnostic groups and thus do not seem to be directly affected by presence of MCI or AD.

### 3.2. Sex-stratified analyses: substantial differences in the association of AD biomarkers and blood metabolite concentrations between the sexes

To investigate if sex modified the association between AD endophenotypes and metabolite concentrations, we tested for associations of the three A-T-N biomarkers with concentrations of 140 blood metabolites. We did this in the full data set and separately in each sex using multivariable linear and logistic regression, followed by analysis of heterogeneity of effects between sexes. **Table 2** lists the results for all metabolite-phenotype combinations, and analyses of sex-by-metabolite interaction effects on A-T-N biomarkers, that fulfilled at least one of the following criteria: (a) associations significant (Bonferroni threshold of *P* < 9.09 × 10^−4^) in the full cohort; (b) associations Bonferroni-significant in one sex; (c) associations that showed suggestive significance (*P* < 0.05) in one sex coupled with significance for effect heterogeneity between female and male effect estimates. Results for all models are provided in Supplementary Table 2. Systematic comparison of estimated effects in each sex for all metabolites is shown in **Figure 1**. Based on this comparison, we classified metabolite – A-T-N biomarker associations into homogenous effects if metabolites showed very similar effects in their association to the biomarker for both sexes, and heterogeneous effects if effects showed opposite effect directions of the same metabolite for men and women or substantially larger effects in one sex leading to significant heterogeneity and/or sex-metabolite interaction. Effects only significant in one sex with either significant effect heterogeneity between males and females or significant sex-metabolite interaction were considered sex-specific.

**Table 2:**
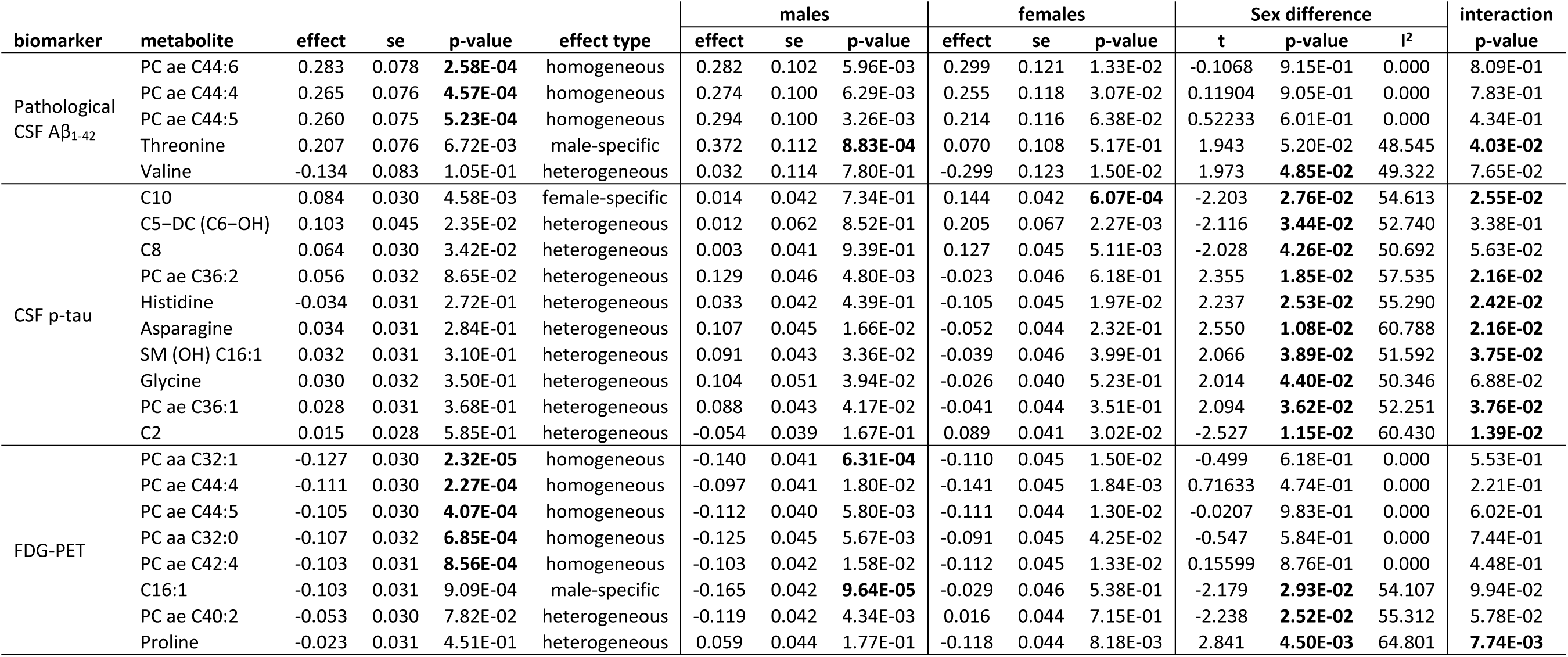
Metabolite associations with A-T-N biomarkers that are either Bonferroni significant in the full sample, one sex, or show nominal significance both in one sex and for effect heterogeneity. Given are regression results for the full sample and both sexes, as well as heterogeneity estimates and the p-value for sex * metabolite interactions.

**Figure 1:** Scatter plots showing Z-scores of effect estimates of metabolite associations with A-T-N biomarkers for males (x-axis) versus females (y-axis). Homogeneous effects (i.e. those with same effect direction and comparable effect size) are located close to the diagonal, heterogeneous effects are located close to the anti-diagonal, and sex-specific effects are located close to the x-axis for male-specific and y-axis for female-specific effects. Homogeneous overall significant results are drawn as diamonds, effects with significant heterogeneity are drawn as rectangles, and effects significant in only one sex are drawn as triangles. Metabolites marked by an asterisk are significant in one sex only and simultaneously show significant heterogeneity. Sex-specificity is further illustrated by a color scale (blue: females; green: males). On the upper right panel, example boxplots of metabolite residuals (obtained by regressing out included covariates) for each effect type are shown separately for the sexes with (in dark red) and without (in light red) CSF Aβ_1-42_ pathology, respectively.

#### 3.2.1. Homogeneous effects

Metabolites with homogenous effects lie on or close to the diagonal going through the first and third quadrant when plotting the effect estimates in women against those in men (**Figure 1**). We identified 8 significant homogenous metabolite-phenotype associations with A-T-N biomarkers: CSF Aβ_1-42_ pathology was significantly associated with levels of 3 related ether-containing PCs (PC ae C44:4, PC ae C44:5, PC ae C44:6). Two (PC ae C44:4, PC ae C44:5) were also significantly associated with brain glucose uptake (FDG-PET) in addition to 3 other PCs (PC aa C32:1, PC aa C32:0, PC ae C42:4). For p-tau, we found no homogeneous, overall significant associations. Notably, none of the homogenous associations showed any indication of effect heterogeneity between sexes, and only one reached significance in the sex-stratified analyses: higher blood levels of the diacyl-PC PC aa C32:1 were associated with lower glucose uptake in brain in the male stratum alone despite lower power.

#### 3.2.2. Heterogeneous effects

Metabolites with heterogeneous effects fall mainly into the second or fourth quadrant (with the exception of sex-specific effects) when contrasting the effect estimates for men and women in the plots for the three A-T-N phenotypes (**Figure 1**). We identified 15 associations in this category (including 3 sex-specific effects). For CSF Aβ_1-42_, we identified 2 heterogeneous effects with threonine showing a sex-specific effect (see paragraph below) with greater effect size in males, and valine with a larger effect in females. While valine was not significantly associated (*P* = 0.78) with CSF Aβ_1-42_ pathology in males, in females, it showed a nominally significant negative association (estimated heterogeneity of *I*^2^ = 49.3%). CSF p-tau had the largest number of heterogeneous associations: acylcarnitines C5-DC (C6-OH), C8, C10 (sex-specific), and C2, and the amino acid histidine showed stronger associations in females, while the related ether-containing PCs PC ae C36:1 and PC ae C36:2, the amino acids asparagine and glycine, and one hydroxy-SM (SM (OH) C16:1) yielded stronger associations in males (all *I*^2^ > 50%). Associations with FDG-PET revealed 3 heterogeneous effects, with ether-containing PC ae C40:2 and the acylcarnitine C16:1 (sex-specific) showing a larger effect in males (*I*^2^ = 55.3%), and proline having a larger effect in females (*I*^2^ = 64.8%). Nine of the 15 reported heterogeneous associations showed opposite effect directions between sexes, and in 7 cases the interaction term (sex * metabolite) was also significantly (at *P* < 0.05) associated with the respective biomarker.

#### 3.2.3. Sex-specific effects

Metabolites with sex-specific effects fall into the area close to the x- (male-specific) or y- (female-specific) axes of the three effect plots for the different A-T-N phenotypes (**Figure 1**). We found three instances of this effect type. Male-specific effects were seen for threonine with pathological CSF Aβ_1-42_ (positive association) and C16:1 with FDG-PET (negative association). One female-specific effect was seen: higher levels of the medium-chain acylcarnitine C10 were associated with higher CSF p-tau. This association was the strongest seen for p-tau in the full cohort analysis, yet seems to be driven by female effects only.

### 3.3. Stratified analyses by *APOE* ε4 status suggest intertwined modulation of metabolite effects by both sex and *APOE* genotype

Prior reports suggested the *APOE* ε4 genotype may exert AD risk predisposition in a sex-dependent way (3, 4, 9, 34). To investigate potential relationships between sex and *APOE* ε4 status on the metabolomic level, we selected the 21 metabolites identified in the previous analyses (**Table 2**) and performed association analyses with the three selected A-T-N biomarkers, now stratified by *APOE* ε4 status and adjusted for sex. Metabolite effects in *APOE* ε4 +/- showed effects from all three of the six-stratified analyses categories (**Table 3**). **Homogenous effects** were noted for the overall significant associations of PC aa C32:1, PC ae C44:4, PC ae C44:5, PC aa C32:0, and PC ae C42:4 with FDG-PET. **Heterogeneous effects** again formed the largest group (n = 11), with proline and glycine showing opposite effect directions on CSF Aβ_1-42_ pathology, and C8, valine, glycine, and proline having opposite effect directions on FDG-PET for *APOE* ε4 +/-, respectively. Five metabolites with heterogeneous effects showed ***APOE* ε4 status-effects**. PC ae C44:6, PC ae C44:4, PC ae C44:5, and PC ae C42:4 showed association with pathological CSF Aβ_1-42_ in *APOE* ε4 carriers; in the case of PC ae C44:6, PC ae C44:5, and PC ae C44:4, the group-specific effects were strong enough to drive the signal to overall significance in the full sample. Also, acylcarnitine C10 showed association with FDG-PET in *APOE* ε4 non-carriers.

**Table 3:**
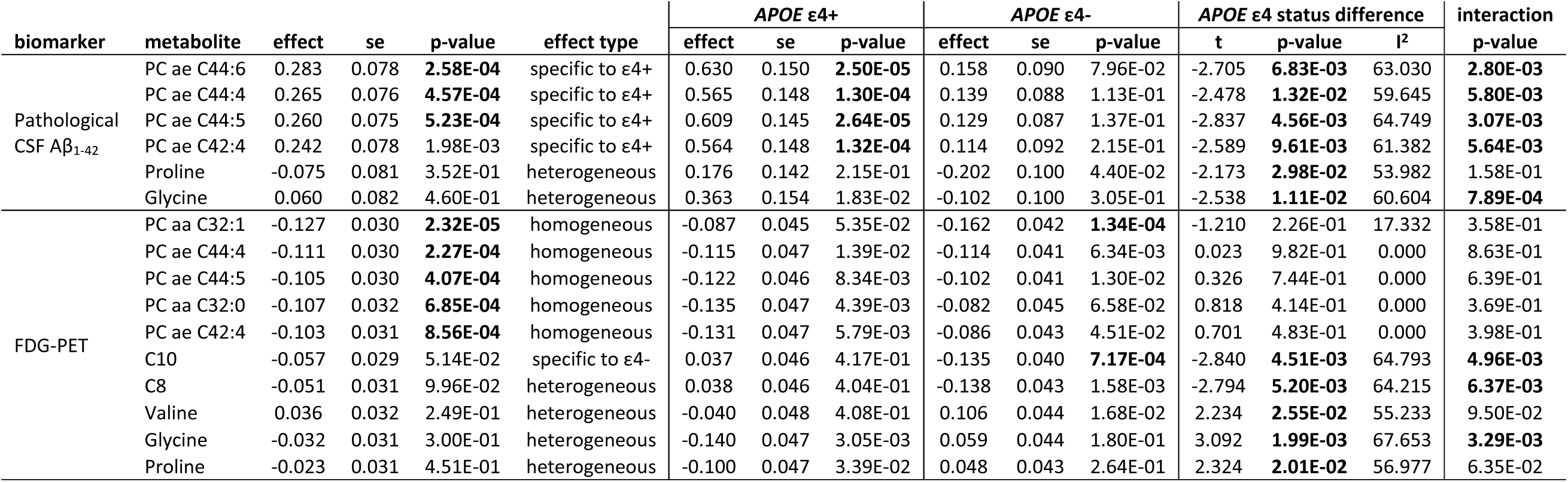
Associations of metabolites identified in the sex-centric analysis with A-T-N biomarkers that are either Bonferroni significant in the full sample, in *APOE* ε4+ or *APOE* ε4-subjects, or show nominal significance both in one *APOE* ε4 status group and for effect heterogeneity. Given are regression results for the full sample and both *APOE* ε4 status groups, as well as heterogeneity estimates and the p-value for *APOE* ε4 status * metabolite interactions.

### 3.4. Combined stratification by sex and *APOE* ε4 status reveals metabolic effects specific to female ε4 carriers

When we stratified separately by sex and *APOE* ε4 status, several metabolites (C8, C10, valine, glycine, proline) showed heterogeneous effects on AD biomarkers in both stratifications. To investigate potential additional subgroup-specific effects, we combined the two stratifications and investigated the selected metabolite set for sex-by-*APOE* ε4 status effect modulations. Although the group of *APOE* ε4-carrying women was the smallest among the strata, all Bonferroni-significant associations were found in this subgroup (**Table 4**): higher levels of three ether-containing PCs (PC ae C42:4, PC ae C44:5, PC ae C44:6) were associated with pathological CSF Aβ_1-42_, higher acylcarnitine C10 was associated with increased CSF p-tau, and higher proline levels were associated with decreased FDG-PET values (**Figure 2**). The latter was not observed in any other performed analyses. Except for the association of C10 with p-tau, we found significant (*P* < 0.05) interaction effects between the metabolites and *APOE* ε4 status on their associated endophenotypes in females only.

**Figure 2:** Boxplots showing residuals of proline levels (derived by regressing out covariate effects) for **A:** the full sample; **B:** 1-fold stratifications by sex; **C:** 1-fold stratification by *APOE* ε4 status; and **D:** 2-fold stratification by both sex and *APOE* ε4 status; separately for high (light blue) and low (darker blue; derived by mean-split) FDG-PET values. The only subgroup showing a significant difference in proline levels are *APOE* ε4+ females with substantially higher levels in subjects with lower brain glucose uptake.

**Table 4:**
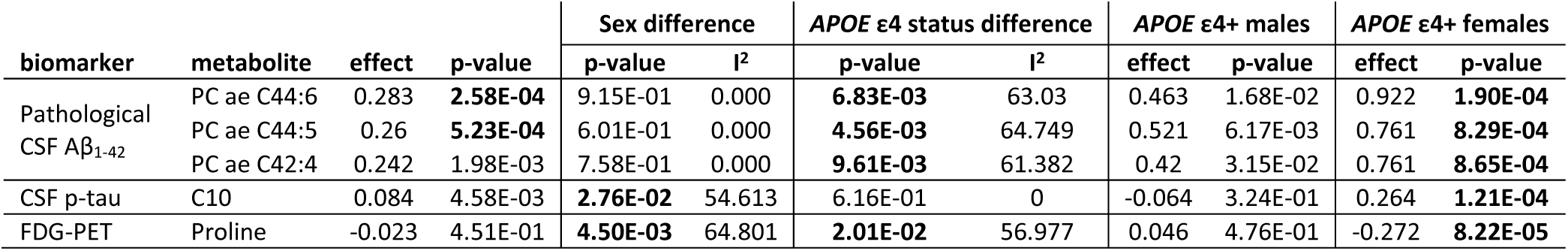
Significant metabolite effects in the combined stratification (sex by *APOE* ε4 status) on A-T-N biomarkers are driven by or limited to *APOE* ε4+ females. Given are regression results for the full sample, *APOE* ε4+ males, *APOE* ε4+ females, as well as heterogeneity estimates by sex and *APOE* ε4 status. The only metabolite showing effect heterogeneity for both stratification variables was proline in its association with FDG-PET values.

| biomarker | metabolite | effect | p-value | Sex difference | | *APOE* ε4 status difference | | *APOE* ε4+ males | | *APOE* ε4+ females | |
| --- | --- | --- | --- | --- | --- | --- | --- | --- | --- | --- | --- |
|  |  |  |  | p-value | I<sup>2</sup> | p-value | I<sup>2</sup> | effect | p-value | effect | p-value |
| Pathological CSF Aβ<sub>1-42</sub> | PC ae C44:6 | 0.283 | **2.58E-04** | 9.15E-01 | 0.000 | **6.83E-03** | 63.03 | 0.463 | 1.68E-02 | 0.922 | **1.90E-04** |
|  | PC ae C44:5 | 0.26 | **5.23E-04** | 6.01E-01 | 0.000 | **4.56E-03** | 64.749 | 0.521 | 6.17E-03 | 0.761 | **8.29E-04** |
|  | PC ae C42:4 | 0.242 | 1.98E-03 | 7.58E-01 | 0.000 | **9.61E-03** | 61.382 | 0.42 | 3.15E-02 | 0.761 | **8.65E-04** |
| CSF p-tau | C10 | 0.084 | 4.58E-03 | **2.76E-02** | 54.613 | 6.16E-01 | 0 | -0.064 | 3.24E-01 | 0.264 | **1.21E-04** |
| FDG-PET | Proline | -0.023 | 4.51E-01 | **4.50E-03** | 64.801 | **2.01E-02** | 56.977 | 0.046 | 4.76E-01 | -0.272 | **8.22E-05** |

## 4. Discussion

We investigated the influence of sex and *APOE* ε4 status on metabolic alterations related to representative A-T-N biomarkers. Using stratified analyses and systematic comparison of the effects estimated for the sexes, we revealed substantial differences between men and women in their associations of blood metabolite levels with these AD biomarkers, although known sexual dimorphisms of metabolite levels themselves were unaffected by AD.

Differences between the sexes were largest for associations of metabolites and CSF p-tau levels. Notably, this biomarker was not significantly associated with any metabolite when including all subjects and adjusting for both sex and *APOE* ε4 status, yet analysis stratified by sex (but still adjusted for *APOE* ε4 status) revealed a significant, female-specific metabolite/CSF p-tau association despite the smaller sample size. In contrast, for CSF Aβ_1-42_ and FDG-PET, in addition to heterogeneous, sex-specific effects, we also found homogenous effects in which metabolite concentrations showed the same trends of metabolite levels correlating with CSF Aβ_1-42_ pathology and/or lower brain glucose uptake in both sexes.

For many of the metabolites with sex-specific effects, we further observed significant effect heterogeneity between *APOE* ε4 carriers and non-carriers, suggesting intertwined modulation of metabolic effects by sex and *APOE* genotype. Indeed, two-fold stratification revealed metabolite associations that were either driven by or specific to the group with presumably highest risk: *APOE* ε4+ females. Thus, our results demonstrate the importance of stratified analyses for getting insights into metabolic underpinnings of AD that are seemingly restricted to a specific patient group.

### 4.1 Metabolic effect heterogeneity suggests sex-specific differences in energy homeostasis, alternative energy sources, and stress response in AD

The metabolites showing effect heterogeneity across AD biomarkers in this study highlight sex-specific dysregulations of **energy metabolism** (acylcarnitines C2, C5-DC/C6-OH, C8, C10, C16:1 for lipid-based energy metabolism (35); amino acids valine, glycine, proline as markers for glucogenic and ketogenic energy metabolism (36–38)), **energy homeostasis** (asparagine, glycine, proline, histidine (37–40)), and (metabolic/nutrient) **stress response** (threonine, proline, histidine (38, 40)). While these pathways have been linked to AD before, our work presents first evidence and molecular readouts for sex-related metabolic differences in AD.

For instance, our previous report discussed the implication of failing lipid energy metabolism in the context of AD biomarker profiles, starting at the stage of pathological changes in CSF tau levels (20). The current study provides further insights, showing this finding to be predominant in females. More specifically, we observed a significant female-specific association of higher levels of acylcarnitine C10 with increased levels of CSF p-tau, with two other metabolites of this pathway (C8 and C5-DC/C6-OH) narrowly falling short of meeting the Bonferroni threshold. This indicates a sex-specific buildup of medium-chain fatty acids in females, suggesting increased energy demands coupled with impaired energy production via mitochondrial beta-oxidation (35).

Interestingly, the significant heterogeneity of association shown by higher levels of glycine linked to higher CSF p-tau levels in men indicates that energy demands are equally upregulated in males as in females. However, men appear to compensate for this demand by upregulation of glucose energy metabolism, as glycine is a positive marker of active glucose metabolism and insulin sensitivity (37). Findings for acylcarnitines in females are further contrasted by the observed male-specific association of higher levels of the long-chain acylcarnitine C16:1 with decreased brain glucose uptake, which may indicate that in males there is a switch to provision of fatty acids as alternative fuel when glucose-based energy metabolism is less effective. As we did not observe the buildup of medium- and short-chain acylcarnitines as seen in females, we assume that in males, energy production via mitochondrial beta-oxidation may be sustained, at least in early disease.

Evidence corroborating sex-specific processes in energy homeostasis linked to changes in CSF p-tau levels is provided by the significantly lower levels of histidine linked to higher levels of CSF p-tau in women. Depletion of histidine is associated with insulin resistance, inflammatory processes, and oxidative stress, especially in women with metabolic dysregulation (39, 40).

We further identified a heterogeneous association of valine with Aβ_1-42_ pathology, with lower levels in females (*P* < 0.05) but not males. Valine, a BCAA and important energy carrying molecule, is associated with cognitive decline and brain atrophy in AD, and with risk for incident dementia (17, 20). The lower levels observed in AD are in contrast to other complex phenotypes such as type 2 diabetes, insulin resistance, or obesity (36), in which higher levels of BCAAs are found, and may indicate a switch to increased energy consumption via degradation of amino acids in AD. A recent study showed decreasing levels of valine were significantly associated with all-cause mortality (41). Besides implications for energy metabolism, results from our study may thus characterize lower levels of valine also as a marker for increased female vulnerability to pathogenic processes in general and to β-amyloidosis in AD in particular.

### 4.2 Complex interplay between sex, *APOE* ε4 status, and metabolism

The higher effect size of genetic risk for AD exerted by the *APOE* ε4 allele in females compared to males still awaits molecular elucidation. Here we tried to elaborate on potential interrelated risk predispositions from a metabolomic point of view. We therefore investigated whether *APOE* ε4 status may also modulate metabolic readouts of AD-linked A-T-N biomarker profiles identified in sex-centered analyses. Indeed, the majority (68.8%) of observed associations between metabolites and AD biomarkers showed significant heterogeneity between *APOE* ε4 status groups.

The full set of metabolites yielding significant effect heterogeneity when comparing *APOE* ε4-carriers vs. non-carriers (C8, C10, glycine, proline, valine) showed significant heterogeneity estimates in the sex-stratified analyses. We therefore applied two-fold stratification by sex and *APOE* ε4 status to identify potential interactions between both variables (**Supplementary Figure 3**). Several associations showed Bonferroni significance in the group with presumably the highest AD risk: *APOE* ε4+ females. One significant association—higher proline levels with reduced brain glucose uptake—was not observed in the three other strata, the one-fold stratifications, or the full sample, emphasizing the value of more fine-granular stratified analyses as proposed here.

### 4.3 Homogeneous effects seem to represent generic metabolic hallmarks in AD

The heterogeneity of metabolite effects we identified might, in part, explain inconsistencies (e.g., (42) vs. (43)) in associations of metabolites and AD reported in other studies (e.g., if sex and *APOE* genotype are distributed differently and sample sizes are small). Besides the heterogeneous, sex-specific effects observed for metabolite associations with CSF Aβ_1-42_ and FDG-PET biomarkers, we also found associations of these biomarkers with metabolites that showed the same effects in both sexes. In particular, PCs that presumably contain two long-chain fatty acids with, in total, 4 or 5 double bonds (PC ae C44:4, PC ae C44:5) were significant for both AD biomarkers. Such homogeneous metabolite associations should replicate well across studies.

To test this assumption in an independent sample, we performed a targeted analysis using the three PCs associated with CSF Aβ_1-42_ pathology in 86 serum samples from the ROS/MAP cohorts (**Supplementary Text 2**). All three associations were Bonferroni significant (PC ae C44:4 – *P* = 3.73 × 10^−3^; PC ae C44:5 – *P* = 1.15 × 10^−2^; PC ae C44:6 – *P* = 3.28 × 10^−3^) in ROS/MAP with consistent effect directions. In ROS/MAP, we used a different measure of amyloid pathology (total amyloid load in the brain), which is inversely correlated with CSF Aβ_1-42_ levels (44). This inverse relationship was mirrored by metabolite effect estimates. These results provide evidence for homogeneous associations to be relevant across cohorts.

### 4.4 Limitations

First, the reported findings are observational and do not allow for direct causal conclusions. Second, the reported heterogeneity estimates still await replication in an independent cohort with sample sizes appropriate for stratification as well as metabolomics and endophenotypic data available (available ROS/MAP sample sizes were too small to be sufficiently powered). Third, stratified analyses in combination with heterogeneity estimates may identify spurious associations, primarily due to the limited power resulting from group separation. However, we showed that for the majority of the non-homogeneous findings reported (60%), the interaction terms between metabolite levels and sex were also significant in the pooled analysis. When stratifying by *APOE* ε4 status, this was true for an even higher fraction of cases (72.7%). This provides an additional line of support for our conclusions. Finally, we only looked at two risk factors, but others (e.g., type 2 diabetes, high blood pressure) may have metabolic aspects and may reveal even greater molecular heterogeneity.

### 4.5 Conclusion

Effect heterogeneity between subgroups linked to energy metabolism has several important implications for AD research. First, this heterogeneity could explain inconsistencies of metabolomics findings between studies as observed for AD if subjects showed different distributions of variables such as sex and *APOE* ε4 genotype. Second, pooled analysis with model adjustment for such variables as typically applied for sex can mask substantial effects that are relevant for only a subgroup of people. This is also true for combinations of stratifying variables, as we demonstrated for the association of proline with brain glucose uptake in female *APOE* ε4 carriers. Consequently, drug trials may have more success by acknowledging between-group differences and targeting the subgroup with the presumably largest benefit in their inclusion criteria. For energy metabolism, group-specific dietary interventions precisely targeting the respective dysfunctional pathways may pose a promising alternative to *de novo* drug development. Extending our approach by selection of additional variables to further improve stratification may eventually guide the way to personalized medicine.

## Acknowledgements

The results published here are in whole or in part based on data obtained from the AMP-AD Knowledge Portal (doi:10.7303/syn2580853). Data collection and sharing for this project was funded by the Alzheimer’s Disease Neuroimaging Initiative (ADNI) (National Institutes of Health Grant U01 AG024904) and DOD ADNI (Department of Defense award number W81XWH-12-2-0012). ADNI is funded by the National Institute on Aging (NIA), the National Institute of Biomedical Imaging and Bioengineering, and through generous contributions from the following: AbbVie, Alzheimer’s Association; Alzheimer’s Drug Discovery Foundation; Araclon Biotech; BioClinica, Inc.; Biogen; Bristol-Myers Squibb Company; CereSpir, Inc.; Cogstate; Eisai Inc.; Elan Pharmaceuticals, Inc.; Eli Lilly and Company; EuroImmun; F. Hoffmann-La Roche Ltd and its affiliated company Genentech, Inc.; Fujirebio; GE Healthcare; IXICO Ltd.; Janssen Alzheimer Immunotherapy Research & Development, LLC.; Johnson & Johnson Pharmaceutical Research & Development LLC.; Lumosity; Lundbeck; Merck & Co., Inc.; Meso Scale Diagnostics, LLC.; NeuroRx Research; Neurotrack Technologies; Novartis Pharmaceuticals Corporation; Pfizer Inc.; Piramal Imaging; Servier; Takeda Pharmaceutical Company; and Transition Therapeutics. The Canadian Institutes of Health Research is providing funds to support ADNI clinical sites in Canada. Private sector contributions are facilitated by the Foundation for the National Institutes of Health (www.fnih.org). The grantee organization is the Northern California Institute for Research and Education, and the study is coordinated by the Alzheimer’s Therapeutic Research Institute at the University of Southern California. ADNI data are disseminated by the Laboratory for Neuro Imaging at the University of Southern California. Study data provided by the Rush Alzheimer’s Disease Center, Rush University Medical Center, Chicago were supported through funding by NIA grants (P30AG10161, R01AG15819, R01AG17917, R01AG30146, R01AG36836, U01AG32984, U01AG46152), the Illinois Department of Public Health, and the Translational Genomics Research Institute. NIA supported this work (R01 AG059093) and also supported the AD Metabolomics Consortium which is a part of NIA’s national initiatives AMP-AD and M^2^OVE-AD (R01 AG046171, RF1 AG051550, and 3U01 AG024904-09S4). MA, RKD, and GK are supported by NIA grants RF1 AG058942 and R01 AG057452. MA and GK are also supported by the Qatar National Research Fund NPRP8-061-3-011. KN is supported by NIH grants NLM R01 LM012535 and NIA R03 AG054936. PMD is supported by the NIH, Cure Alzheimer’s Fund and the Karen L. Wrenn Trust. PMD has served as an advisor to and or received grants from companies for other projects. PMD also owns shares or serves on the board of companies whose products are not discussed here. PMD is also a co-inventor on patents in this field through Duke University which are unlicensed. RKD is an inventor on key patents in the field of metabolomics including applications for AD. RKD holds equity in Metabolon, a metabolomic company, whose products were not used in the current analyses.

## Supplementary Text 1–Data Availability

Metabolomics datasets from the Biocrates p180 platform used in the currently analyses for the ADNI-1, AGNI GO/2, and ROSMAP cohorts are available via the Accelerating Medicines Partnership-Alzheimer’s Disease (AMP-AD) Knowledge Portal and can be accessed at http://dx.doi.org/10.7303/syn5592519 (ADNI-1), http://dx.doi.org/10.7303/syn9705278 (ADNI GO-2), and http://dx.doi.org/10.7303/syn10235592 (ROSMAP). The full complement of clinical and demographic data for the ADNI cohorts are hosted on the LONI data sharing platform and can be requested at http://adni.loni.usc.edu/data-samples/access-data/. The full complement of clinical and demographic data for the ROSMAP cohorts are available via the RUSH AD Center Resource hub and can be requested at https://www.radc.rush.edu/.

## Supplementary Text 2–Metabolomics data processing

In brief, raw metabolomics data for 182 metabolites was available for 1,681 serum study samples and, for each plate, 2-3 NIST Standard Reference samples were available. Furthermore, we also had blinded duplicated measurements for 19 samples (ADNI-1) and blinded triplicated measurements for 17 samples (ADNI-GO and −2) distributed across plates. We first excluded 22 metabolites with large numbers of missing values (> 40%). Then, we removed plate batch effects using cross-plate mean normalization using NIST metabolite concentrations. Duplicated and triplicated study samples were then used to calculate the coefficients of variation (exclusion criterion >20%) and intra-class correlation (exclusion criterion <0.65) for each metabolite. We removed 20 metabolites that violated these thresholds. Next, we excluded non-fasting samples (n=108), imputed missing metabolite data using half the value of the lower limit of detection per metabolite and plate, log2-transformed metabolite concentrations, centered and scaled distributions to a mean of zero and unit variance and winsorized single outlying values to 3 standard deviations. We then used the Mahalanobis distance for detection of multivariate subject outliers, applying the critical Chi-square value for *P* < 0.01 and removing 42 subjects. Finally, metabolites were adjusted for significant medication effects using stepwise backwards selection (for details see (1)).

## Supplementary Text 3–Phenotype Definitions

Baseline data on CSF Aβ_1-42_, levels of phosphorylated tau protein in the CSF (p-tau), and brain glucose metabolism measured by [^18^F] fluorodeoxyglucose (FDG)-PET biomarkers for ADNI-1, -GO, and −2 participants was downloaded from the LONI online portal at https://ida.loni.usc.edu/. For CSF biomarker data, we used the dataset generated using the validated and highly automated Roche Elecsys electrochemiluminescence immunoassays (1, 2). For FDG-PET, we used an ROI-based measure of average glucose uptake across the left and right angular, left and right temporal and bilateral posterior cingulate regions derived from preprocessed scans (co-registered, averaged, standardized image and voxel size, uniform resolution) and intensity-normalized using a pons ROI to obtain standard uptake value ratio (SUVR) means (3, 4). The pathological CSF Aβ_1-42_ cut-point (1,073 pg/ml) as reported by the ADNI biomarker core for diagnosis-independent mixture modeling (see http://adni.loni.usc.edu/methods/, accessed Oct 2017) was used for categorization since CSF Aβ_1-42_ concentrations were not normally distributed. Processed FDG-PET values were scaled and centered to zero mean and unit variance prior to association analysis; p-tau levels were additionally log2-transformed.

## Supplementary Text 4–Replication Analysis in ROS/MAP

To replicate a subset of the findings reported in this manuscript in an independent cohort, we used metabolomics data obtained from pre-mortem serum samples of 86 deceased participants of the Religious Orders Study and the Rush Memory and Aging Project (ROS/MAP), who had agreed to post-mortem neuropathological examinations, using the same metabolomics kit (AbsoluteIDQ-p180). Of the 86 total participants, 24 were females / 62 males; 52 CN / 24 MCI / 7 AD; mean age was 87.77 (± 6.01) years. Metabolomics data processing was performed very similar as for the ADNI, except that we used a pool of study samples randomly injected across plates instead of NIST standard plasma, and median-instead of mean-based quotient batch removal. We then did a targeted analysis to replicate associations of PC ae C44:4, PC ae C44:5, and PC ae C44:6 with Aβ_1-42_ pathology using post-mortem, neuropathology-derived measures of total amyloid load in the brain. This phenotype was transformed to square root values to get values closer to a normal distribution. Linear regression models were adjusted for age at blood draw, sex, study cohort (ROS vs. MAP), race, number of copies of APOE ε4, as well as years of education. All three p-values were Bonferroni significant when adjusting for three test (p-value threshold of *P* < 1.667), complete result statistics were:

| biomarker | metabolite | effect | se | p-value |
| --- | --- | --- | --- | --- |
| total | PC ae C44:4 | 0.30741 | 0.10277 | 0.00373 |
| amyloid in | PC ae C44:5 | 0.2656 | 0.10257 | 0.01149 |
| the brain | PC ae C44:6 | 0.30992 | 0.10212 | 0.00328 |

**Supplementary Figure 1:**
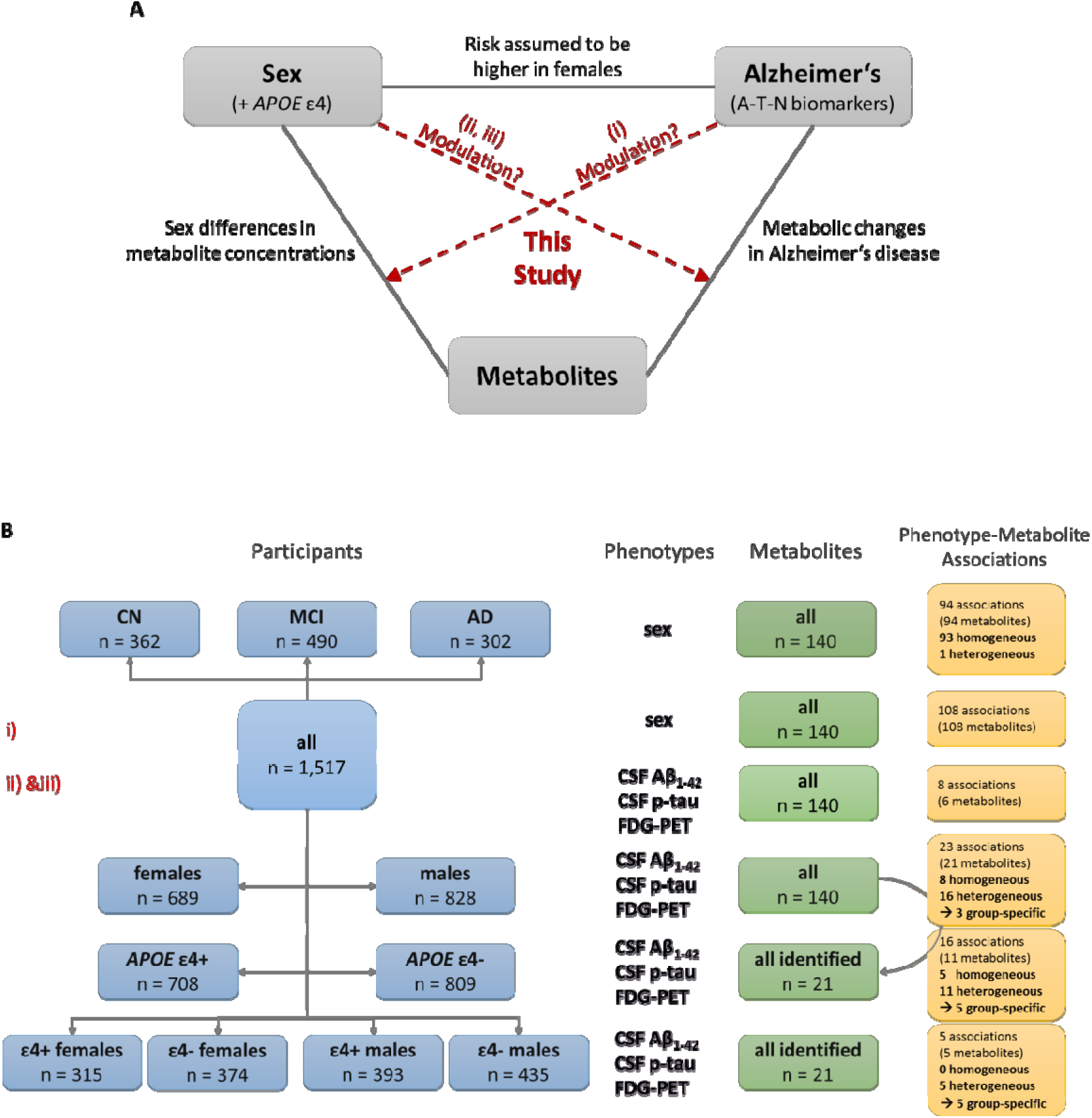
Study rationale and workflow. **A)** This study aims to investigate the relationship between AD, sex, and metabolic readouts in a systematic fashion. The background of this work is: firstly, it has been reported that AD risk may be increased in females; secondly, there are strongly pronounced, highly significant, and often replicated sex differences in metabolite concentrations in the general, healthy population; and, thirdly, we and others have shown that there are significant associations of metabolite levels with AD and its biomarkers. In the current study, we examined: (i) if clinical diagnosis of MCI or AD influences metabolic sex differences as seen in healthy controls, (ii) if sex modulates associations of metabolite levels with three AD biomarkers across the A-T-N spectrum, and, (iii), if effects of metabolites showing sex-based effect heterogeneity in their associations with AD are also modulated by *APOE* ε4 status. **B)** To address the three research questions of this study, we first performed analyses of sex-metabolite associations for 140 metabolites in the ADNI cohort stratified by diagnostic group (question i). Subsequently, we performed phenotype (A/T/N)-metabolite associations for 140 metabolites in the ADNI cohort stratified by sex (question ii) and stratified by *APOE* ε4 status; additionally, we performed phenotype (A/T/N)-metabolite associations for the 21 significantly associated metabolites after stratification by sex plus *APOE* ε4 status (question iii).

**Supplementary Figure 2:**
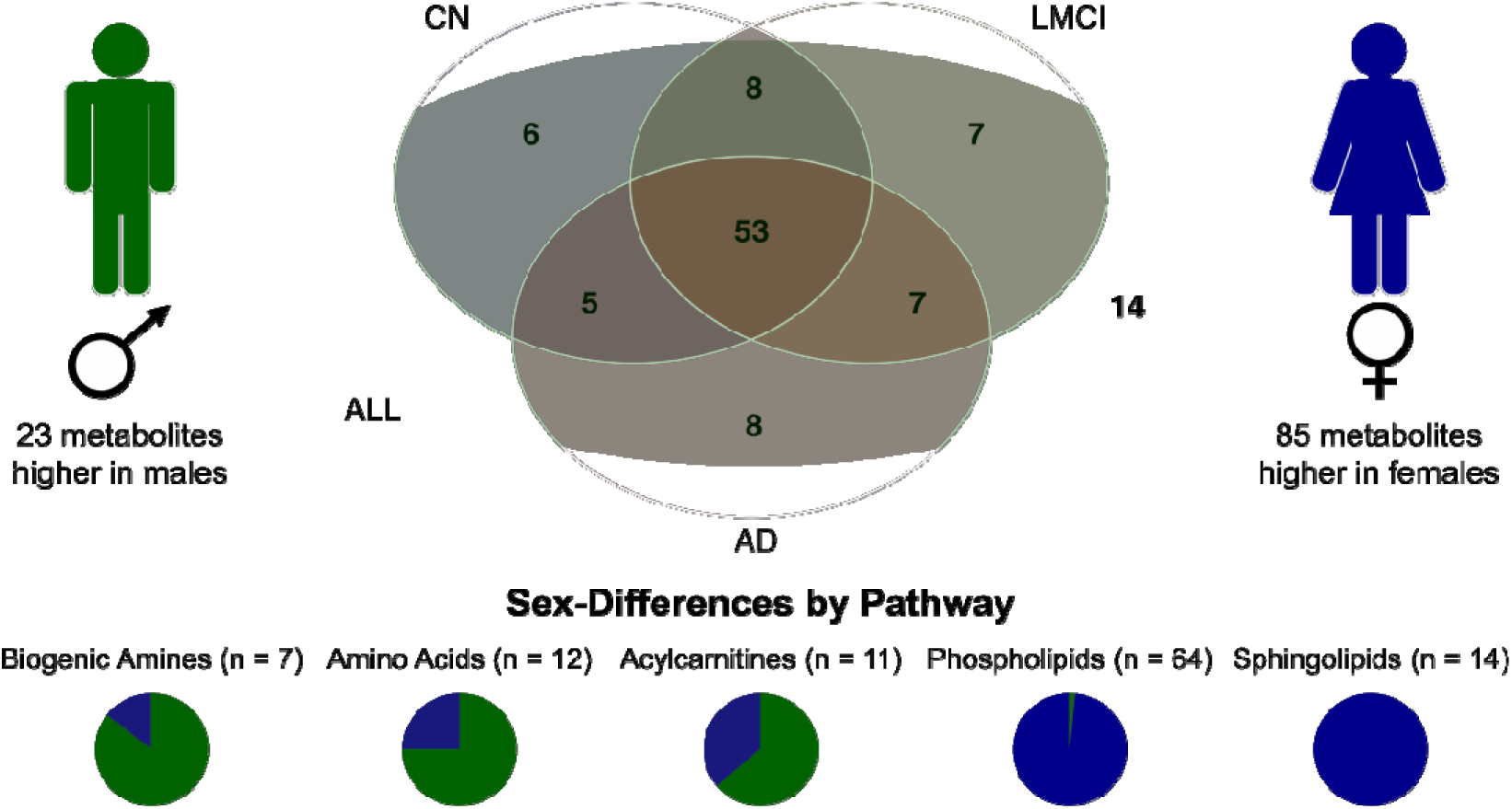
Metabolic sex differences in the ADNI cohorts. We tested whether sex-associated differences in blood metabolite levels differ between patients with probable AD, subjects with MCI, and CN subjects in the ADNI cohorts. We found 108 of 140 metabolites to be significantly associated with sex after multiple testing correction while adjusting for age, BMI, ADNI study phase, and diagnostic group. 70 of these associations replicate previous findings in a healthy population using. All SMs and the majority of PCs were more abundant in women. The majority of biogenic amines, amino acids, and acylcarnitines were more abundant in men. Stratifying subjects by diagnostic group revealed that 53 of the 108 metabolites showing significant sex-differences were also significant in each of the three groups (AD, MCI, CN) alone, while 14 metabolites showed no significant difference in any of the groups, probably due to lower statistical power after stratification. No significant sex-differences were found that were not also significant in the unstratified analysis.

**Supplementary Figure 3.**
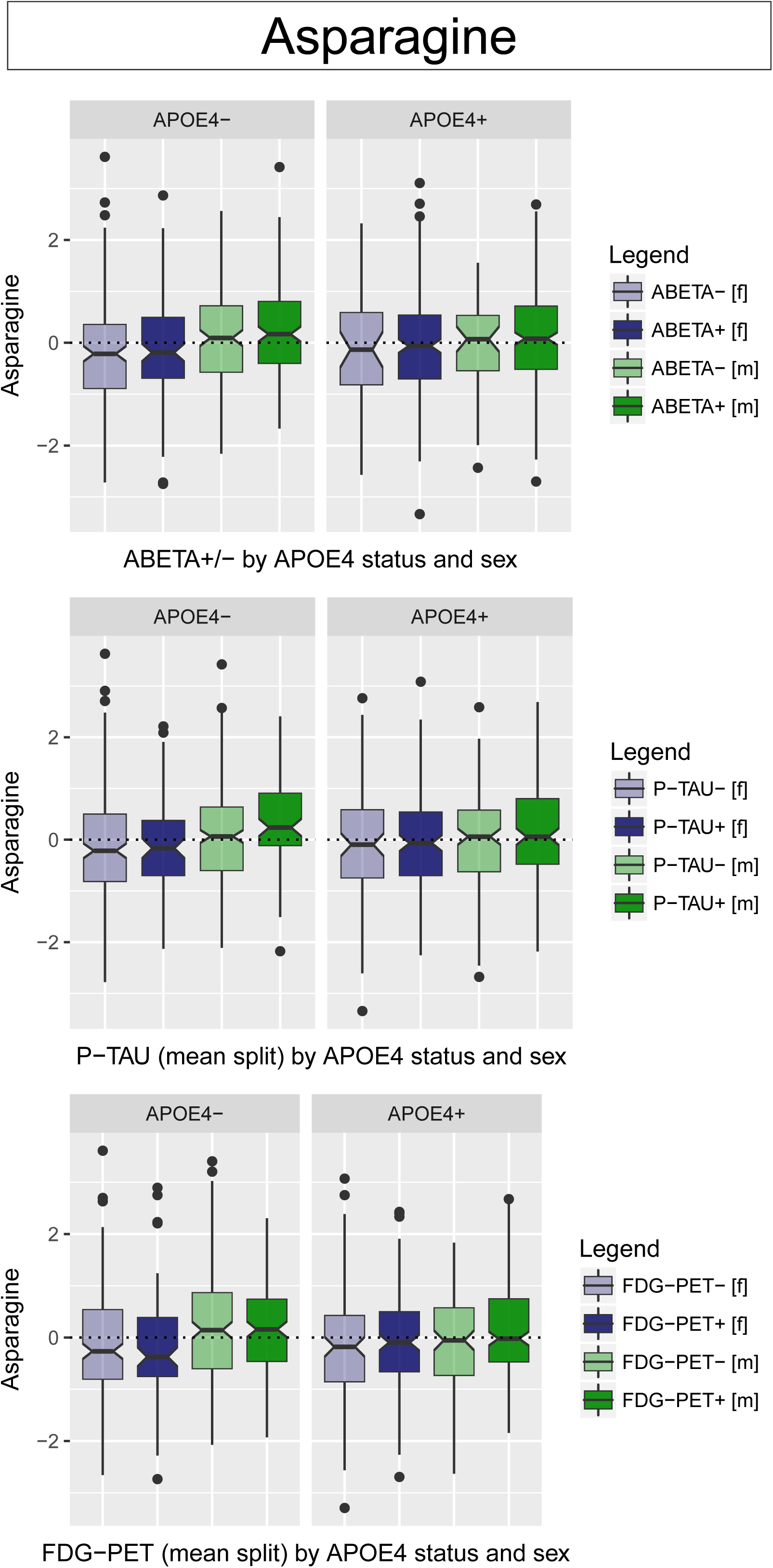
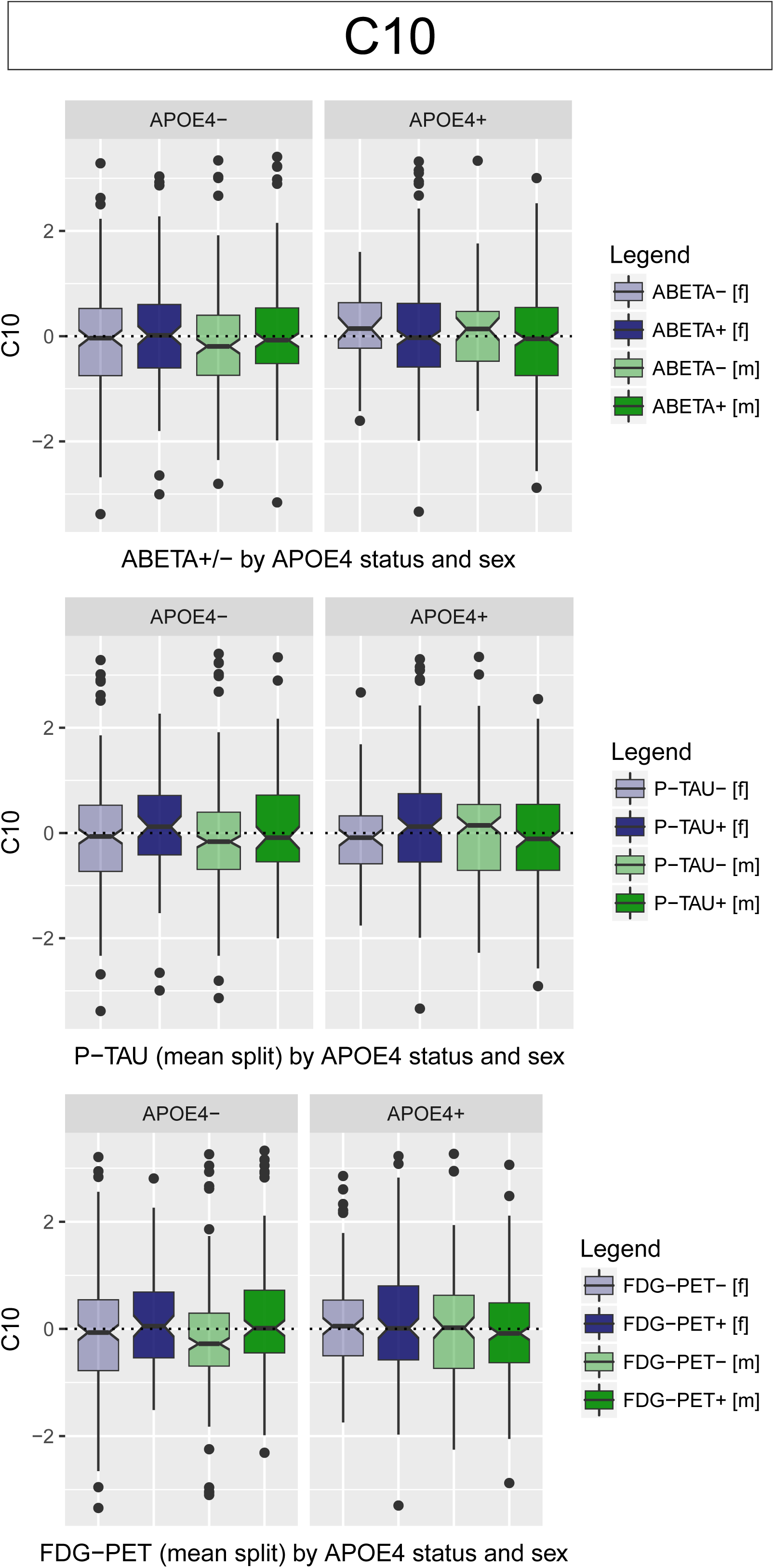
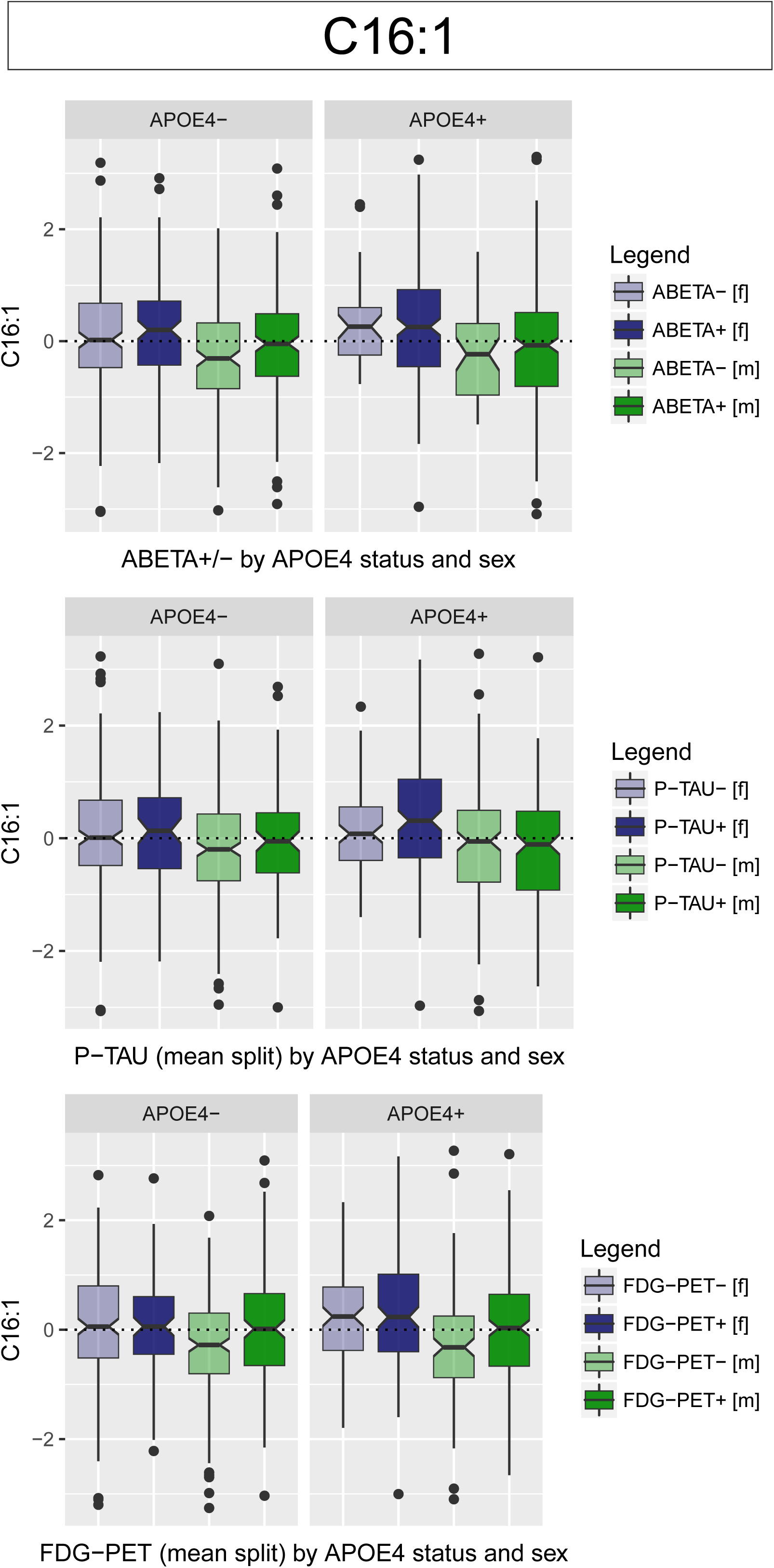
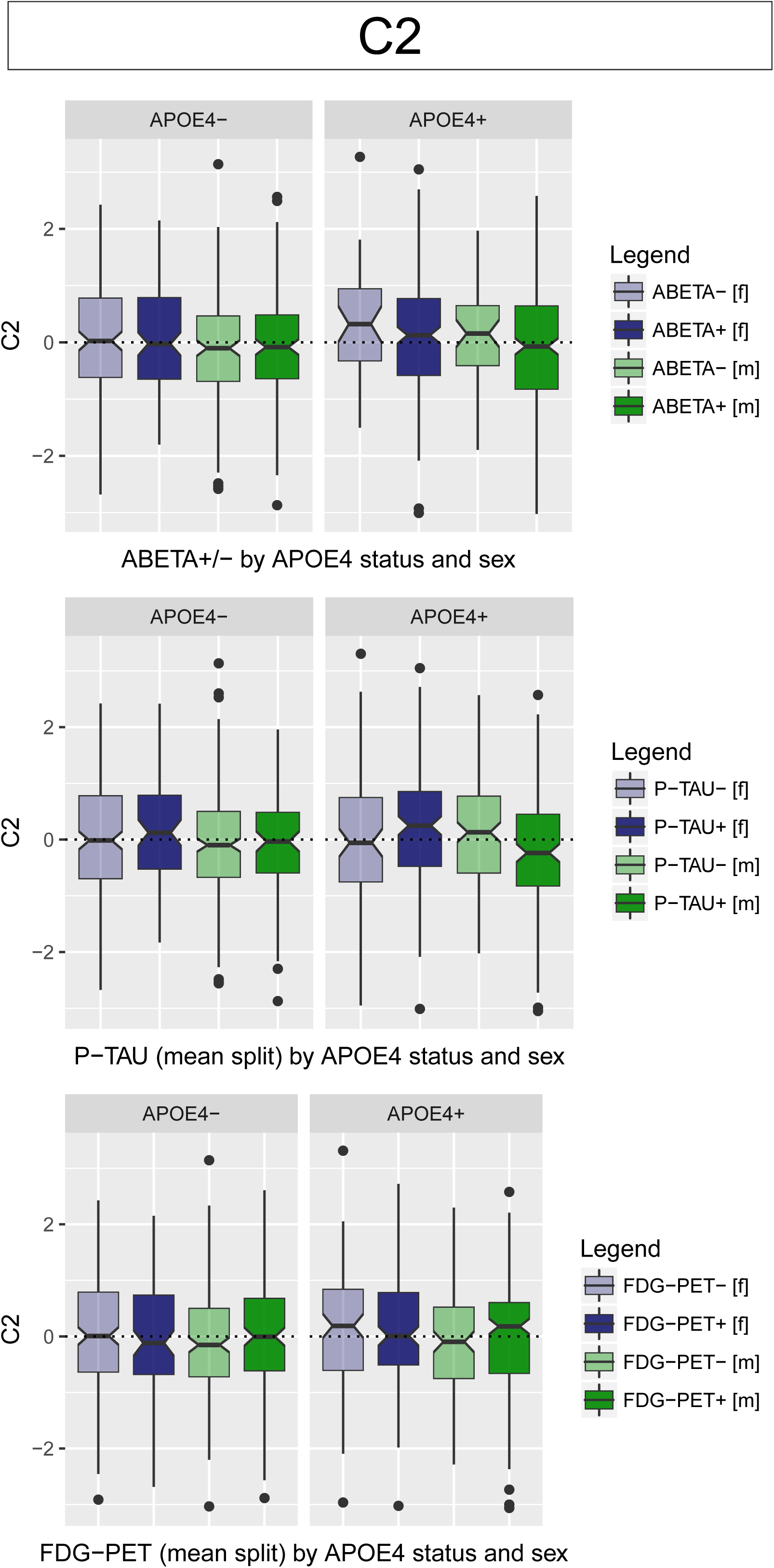
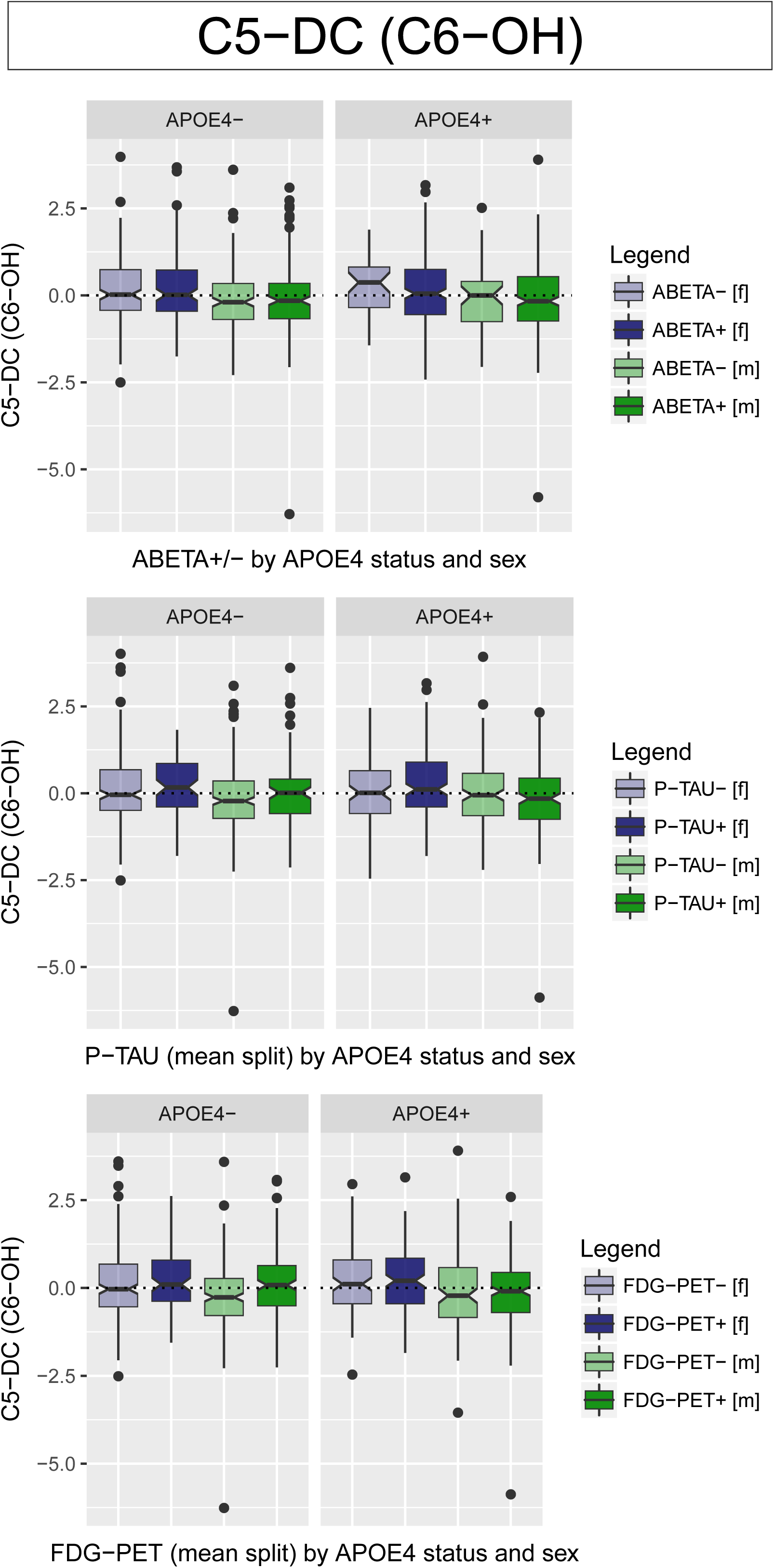
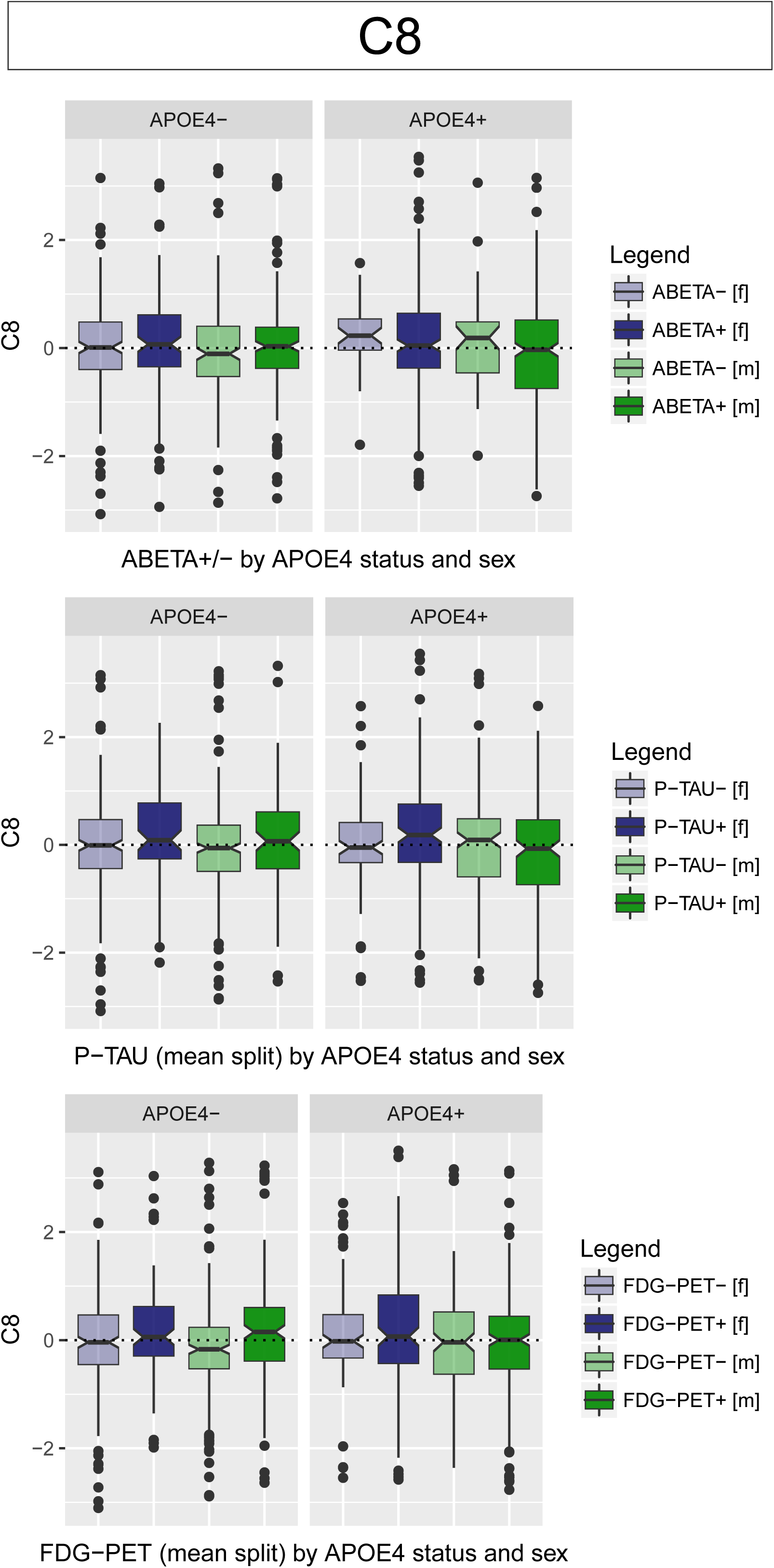
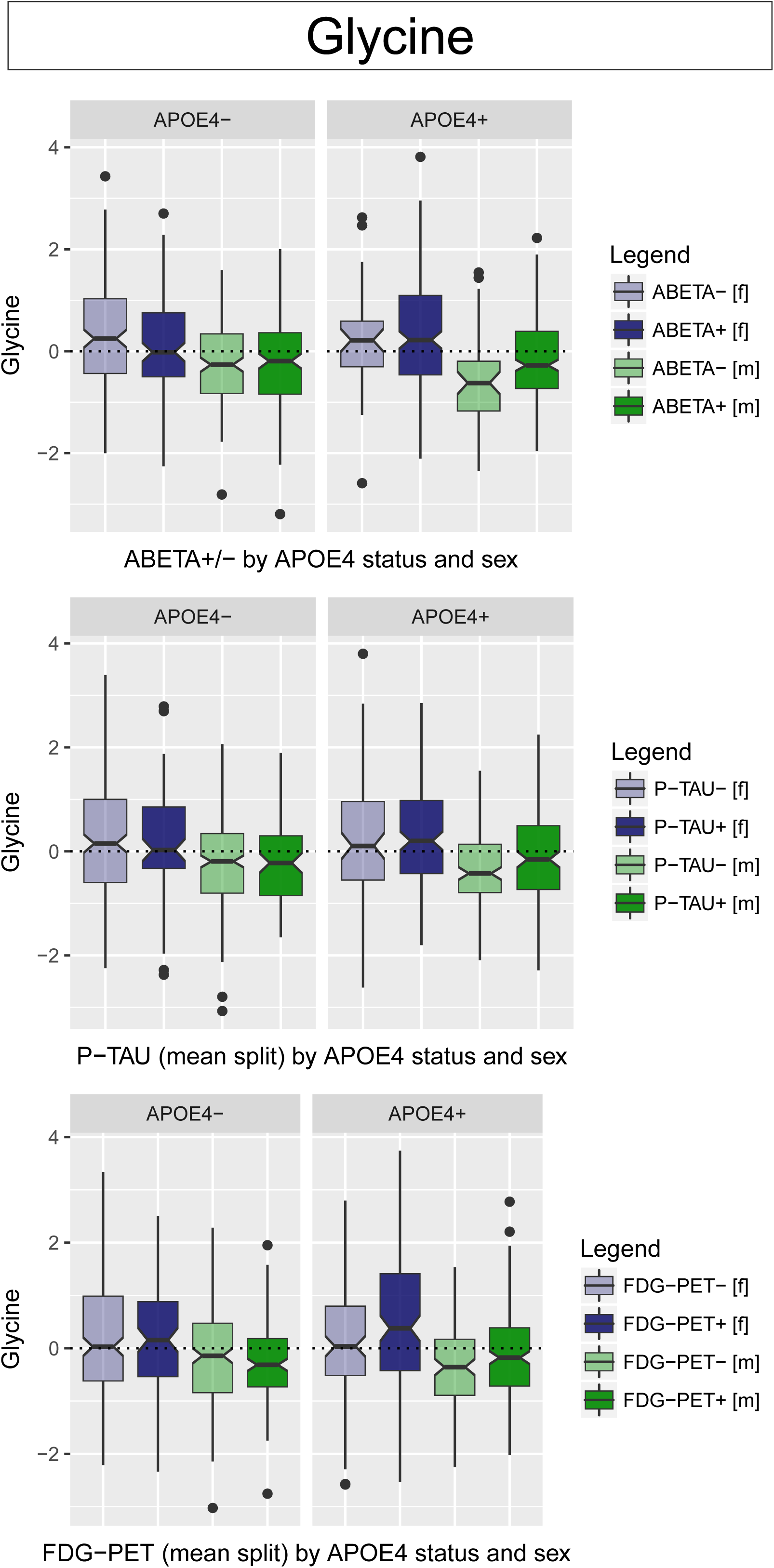
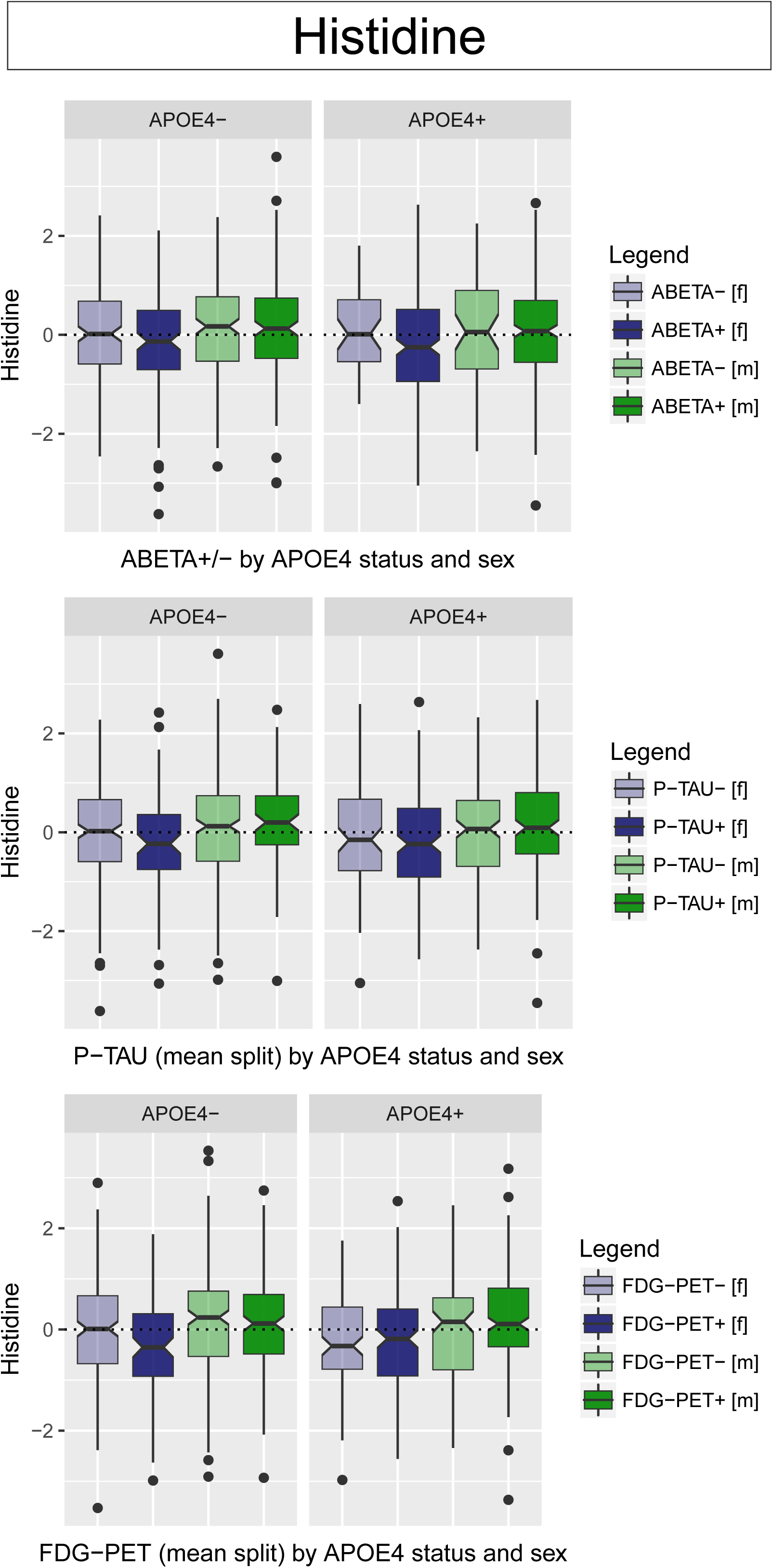
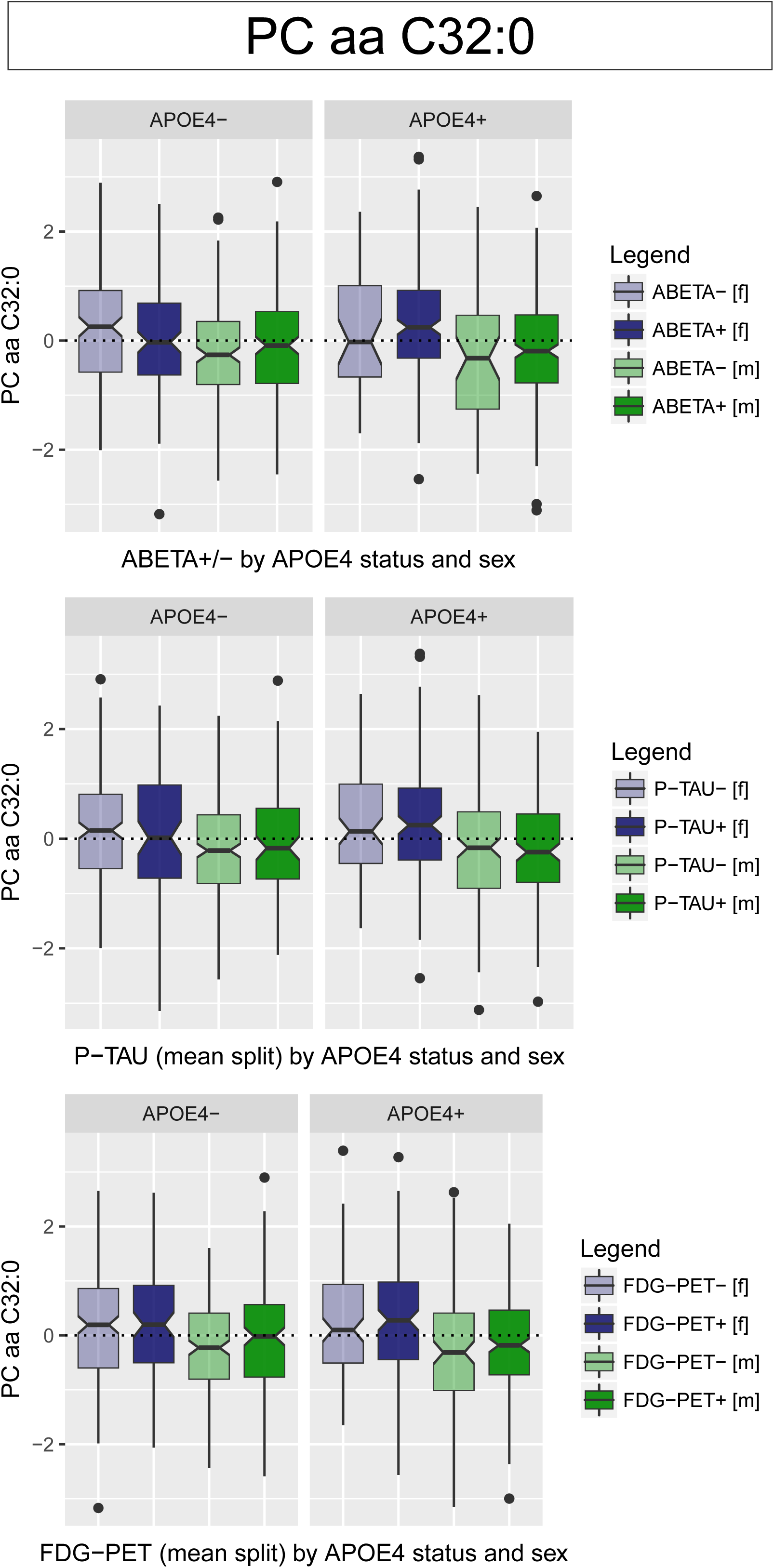
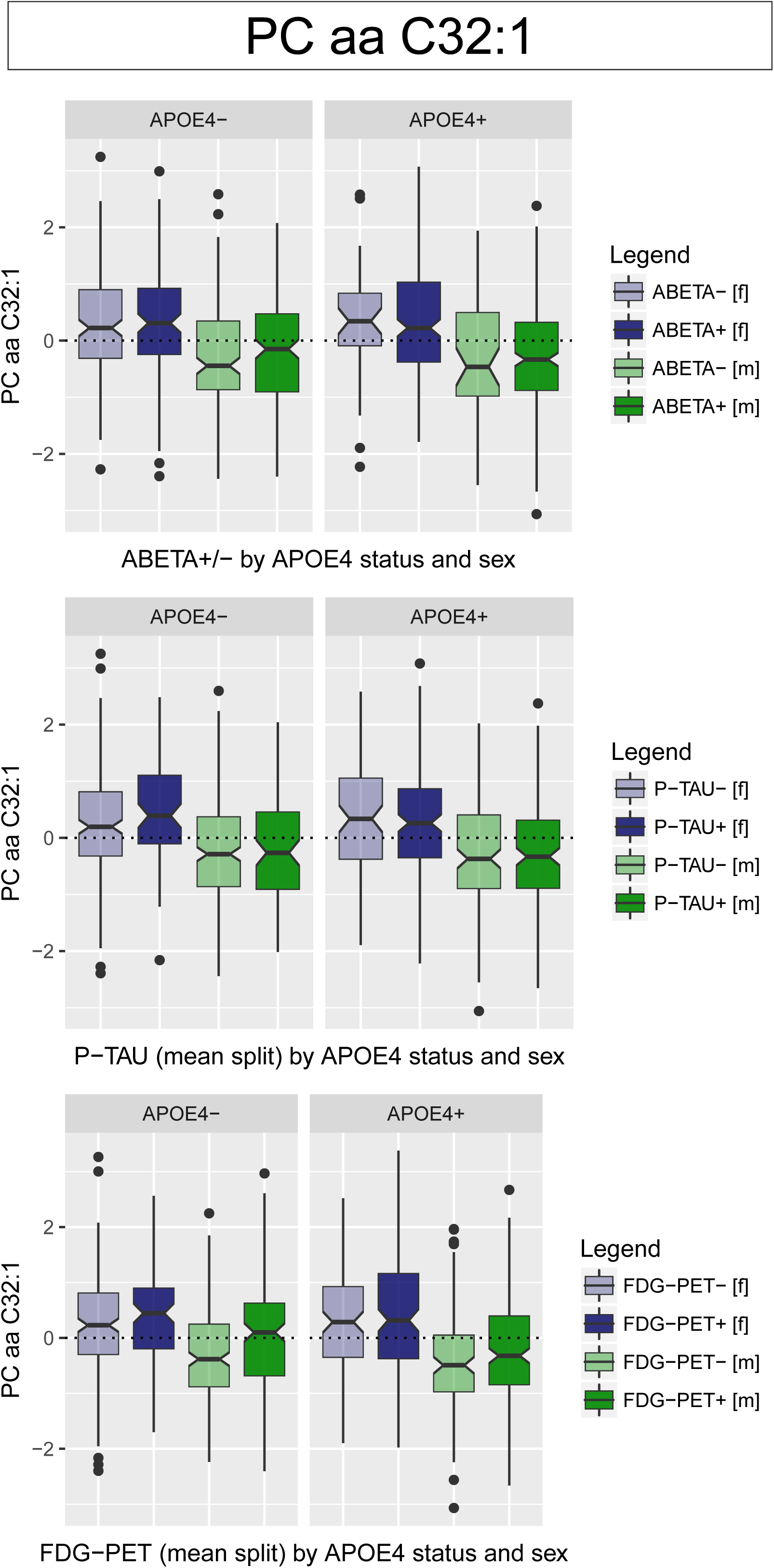
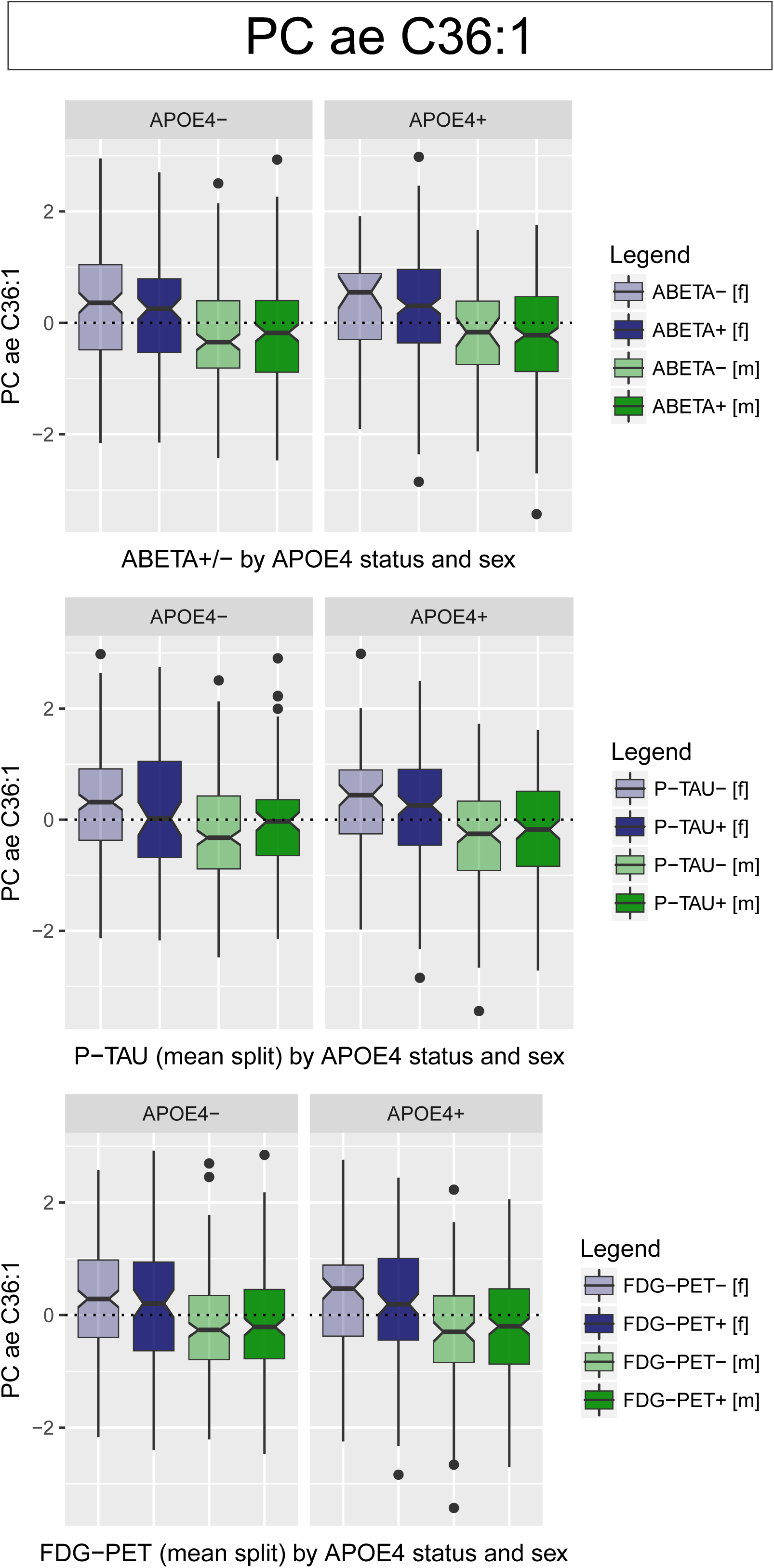
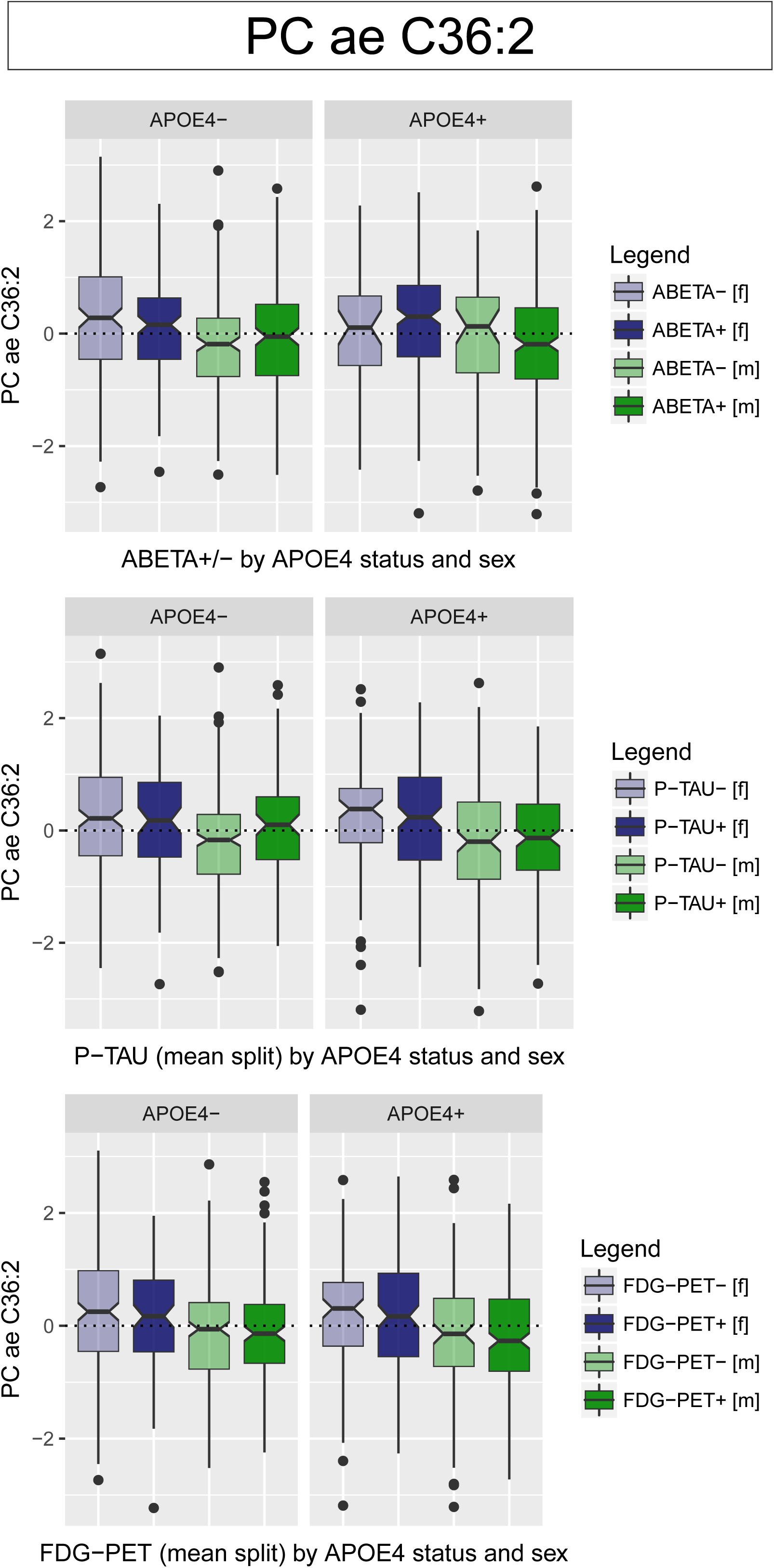
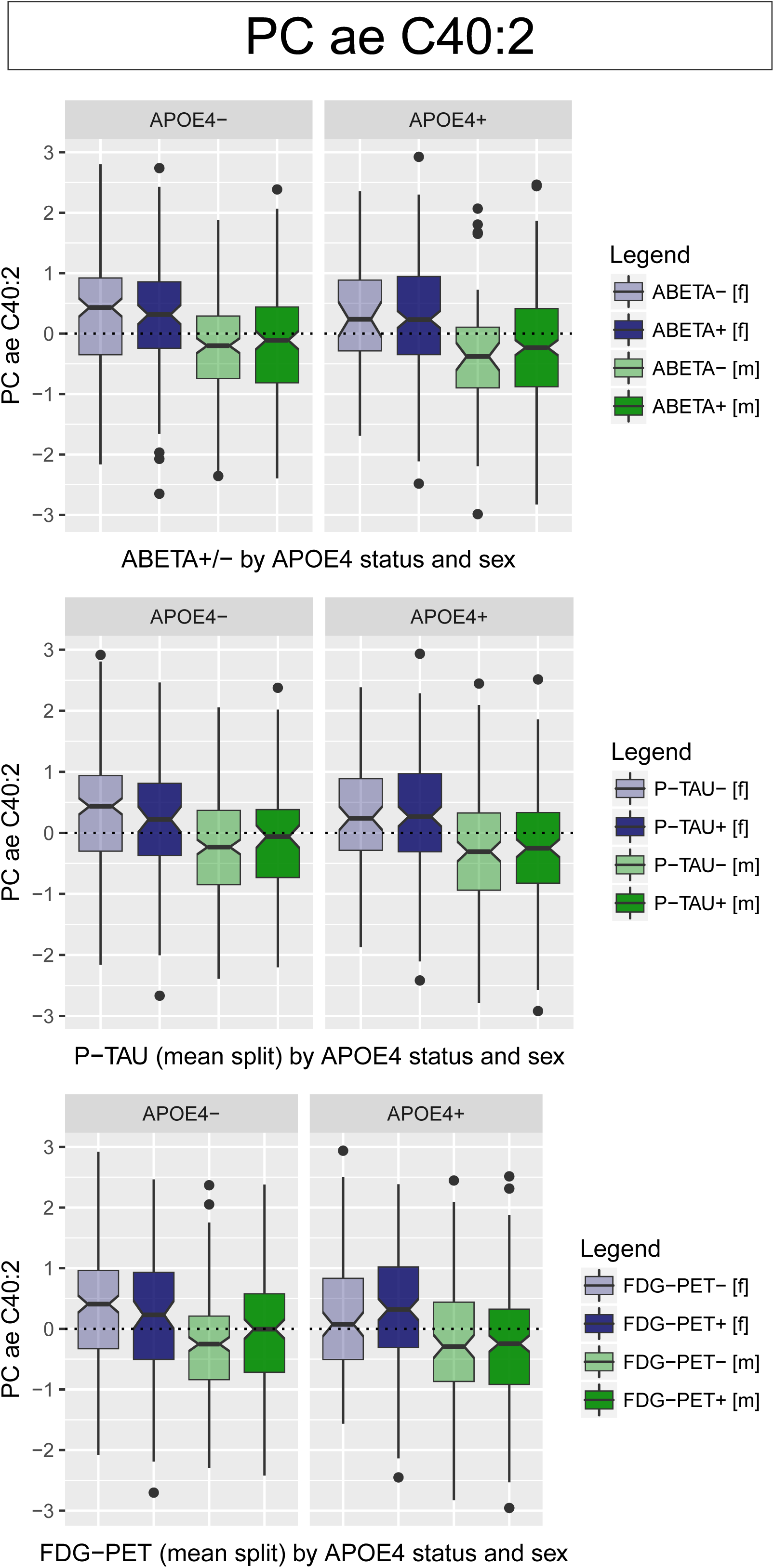
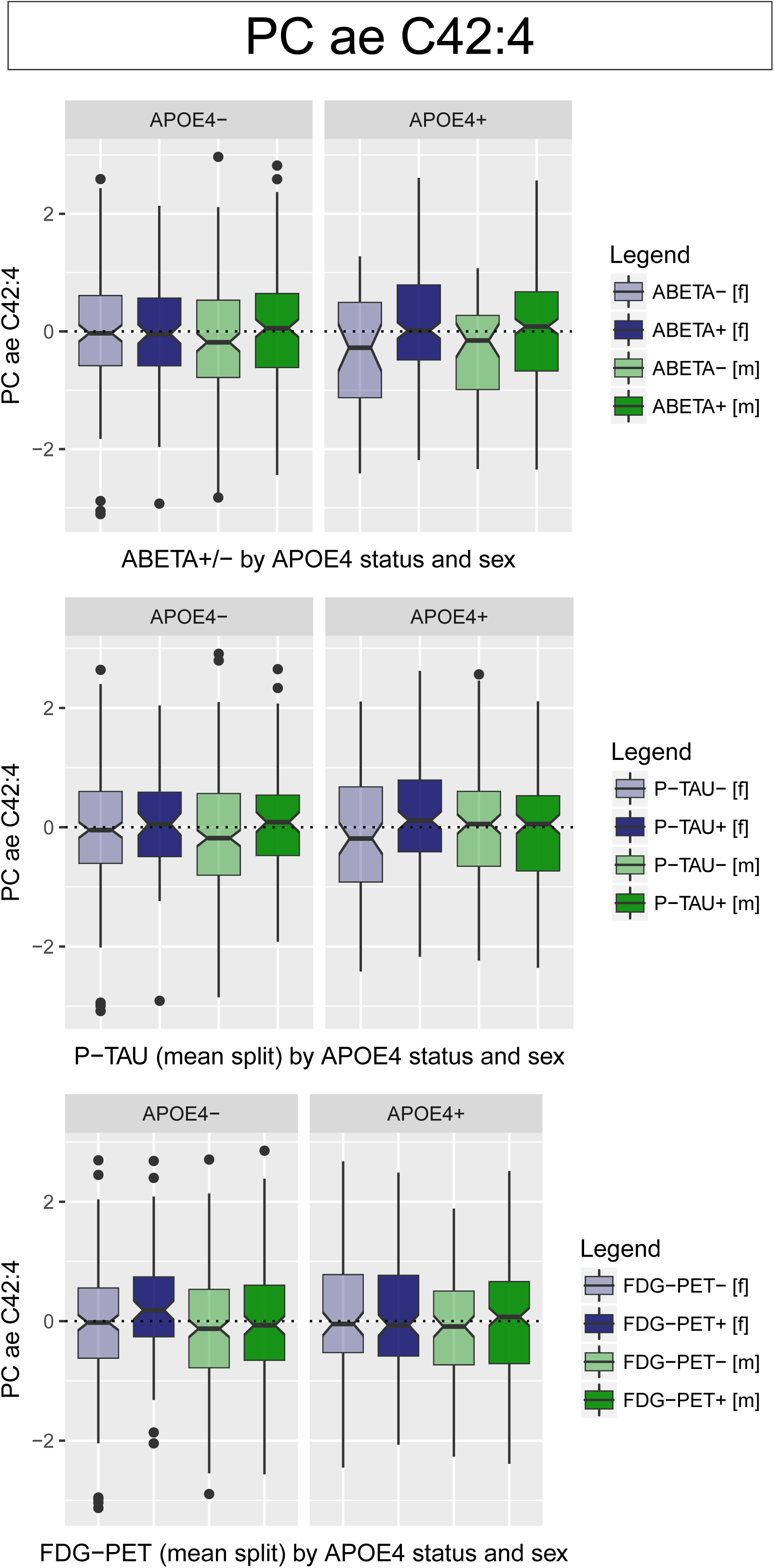
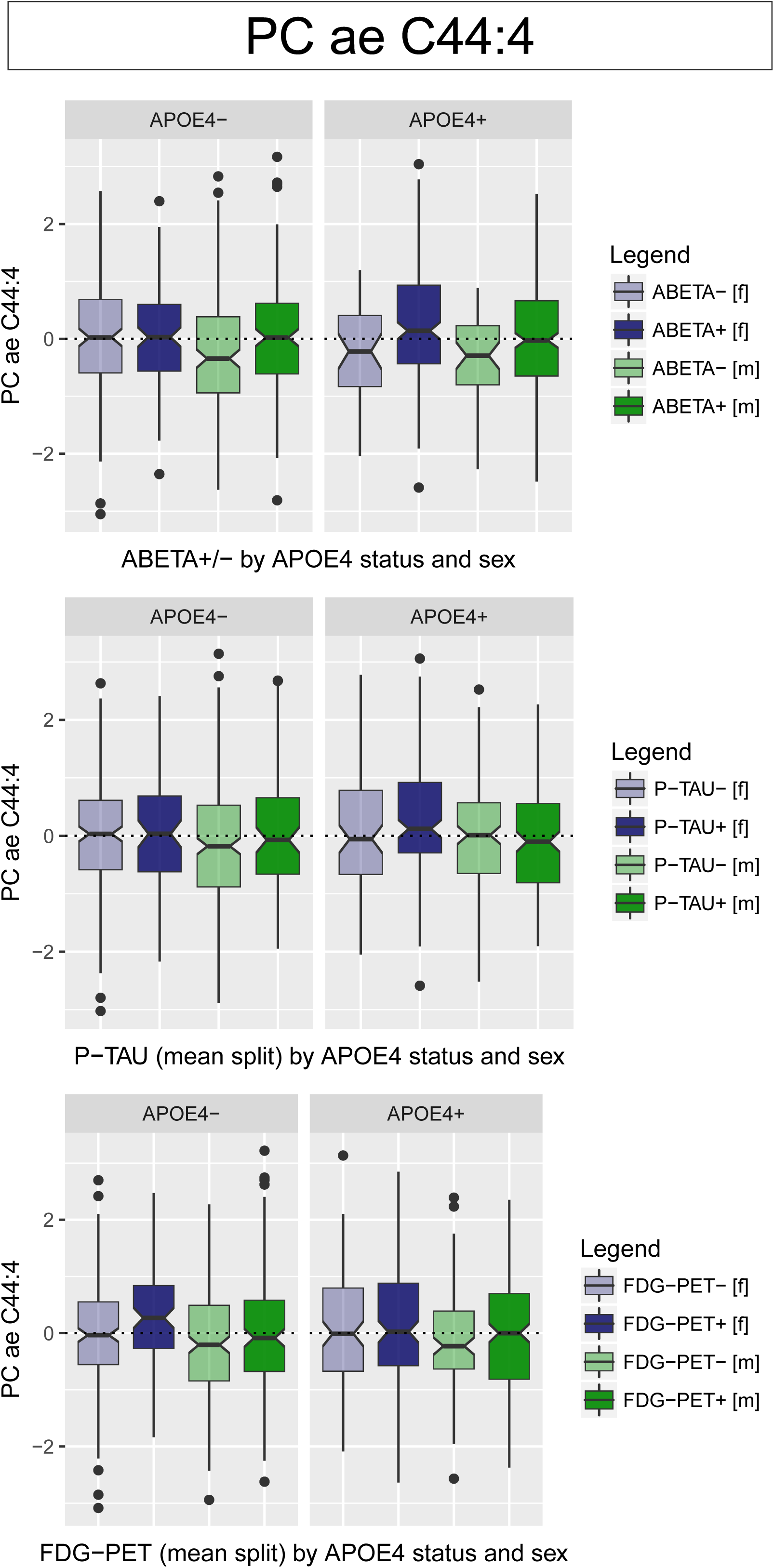
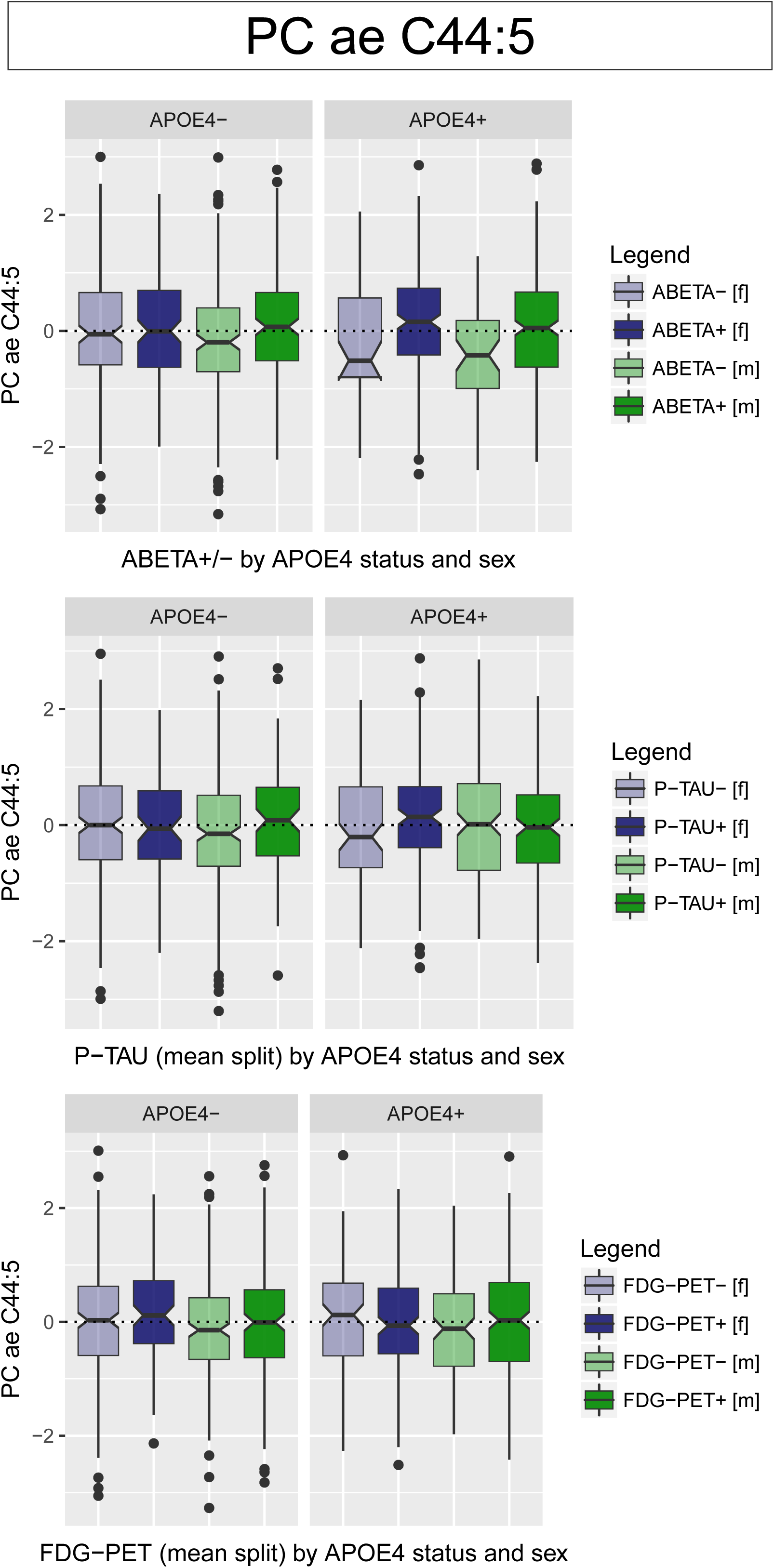
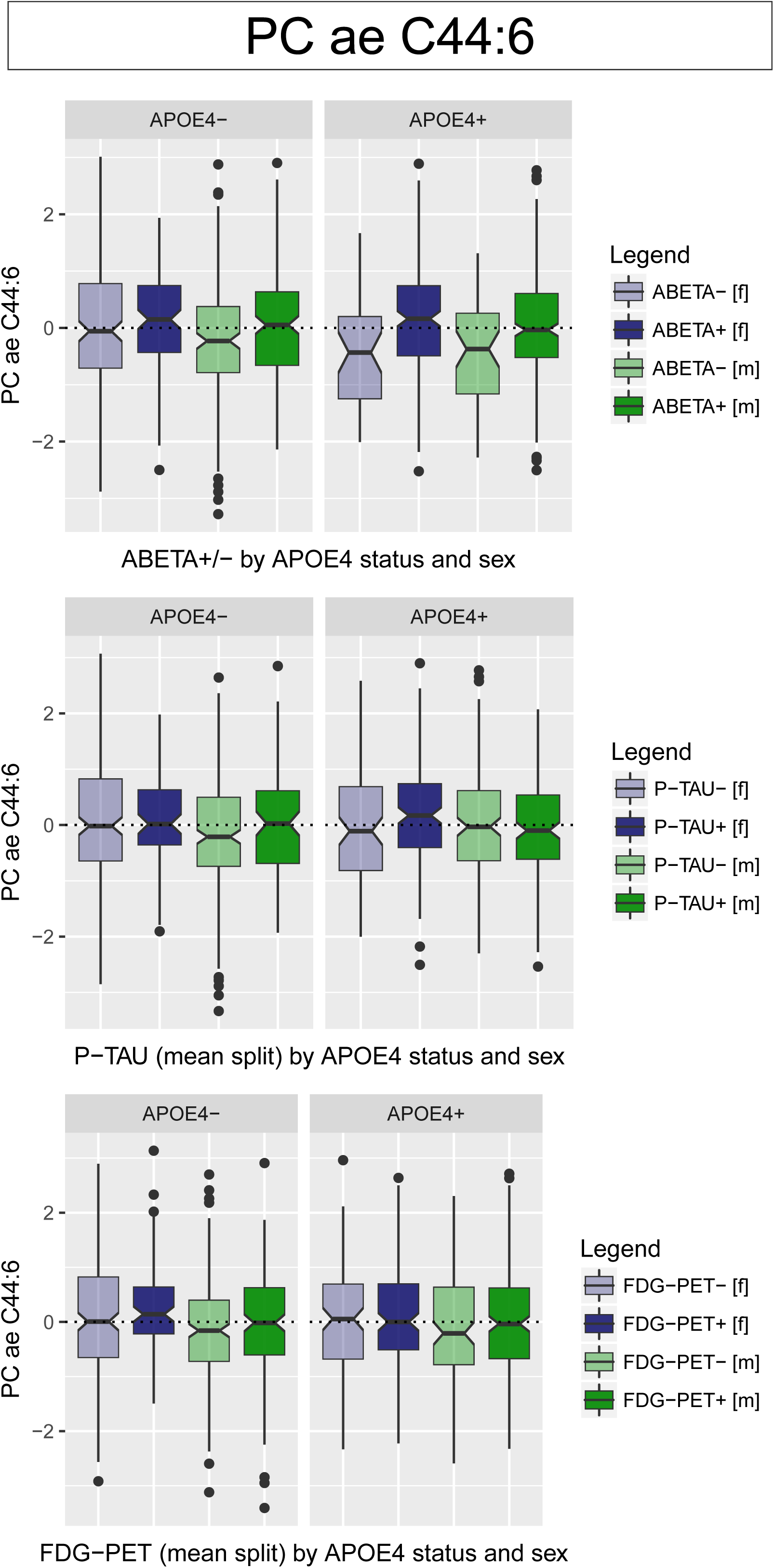
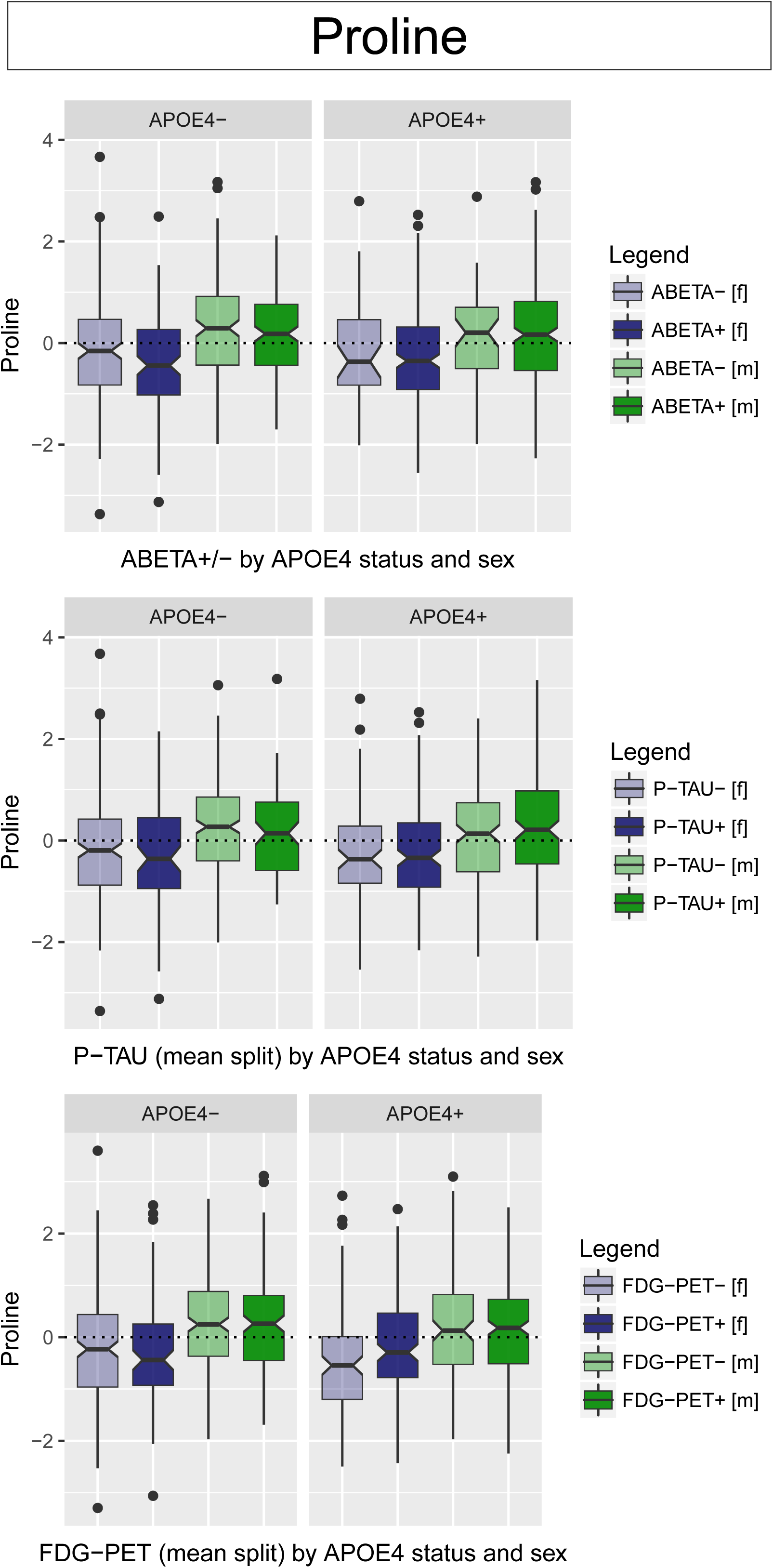
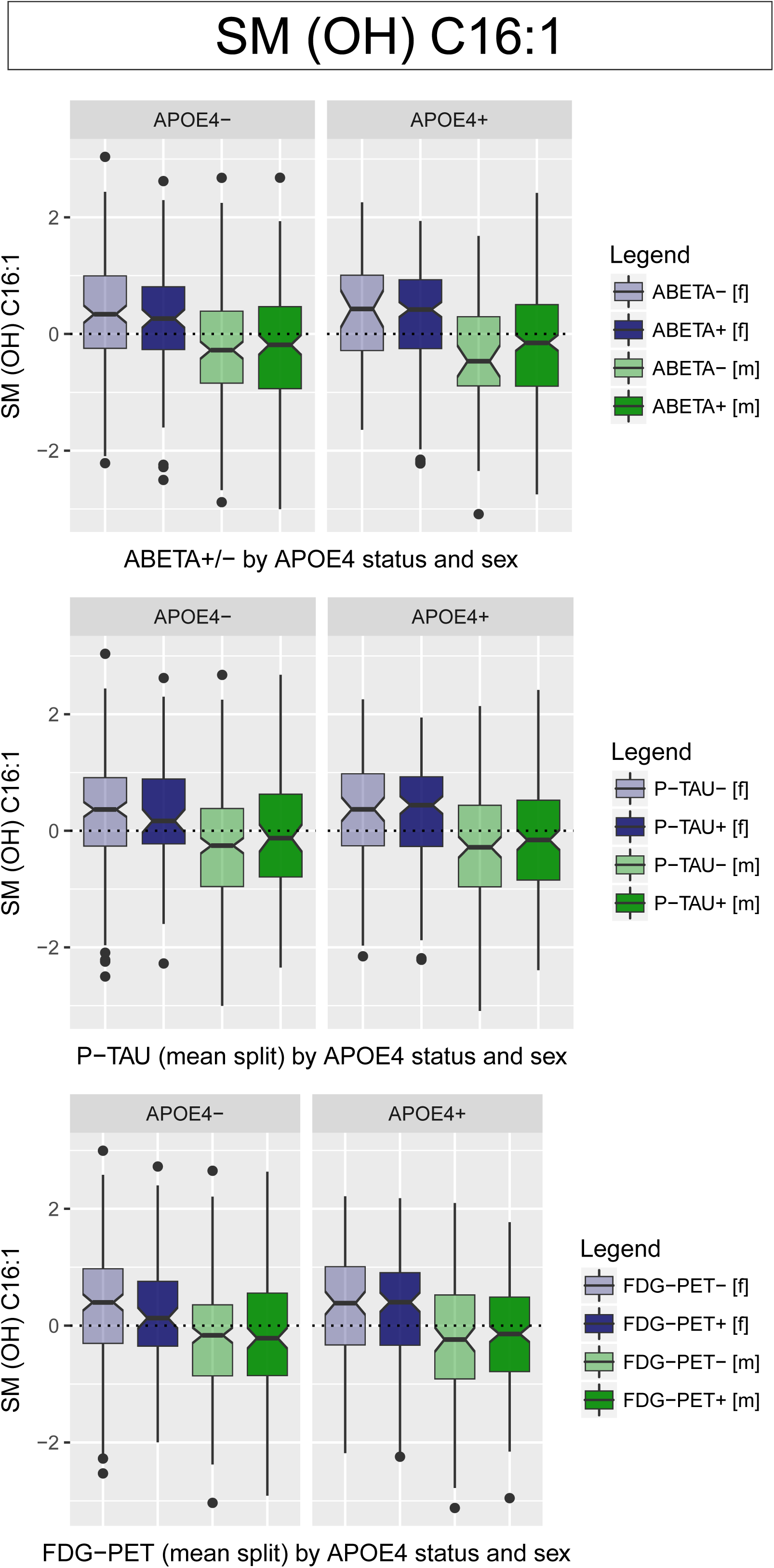
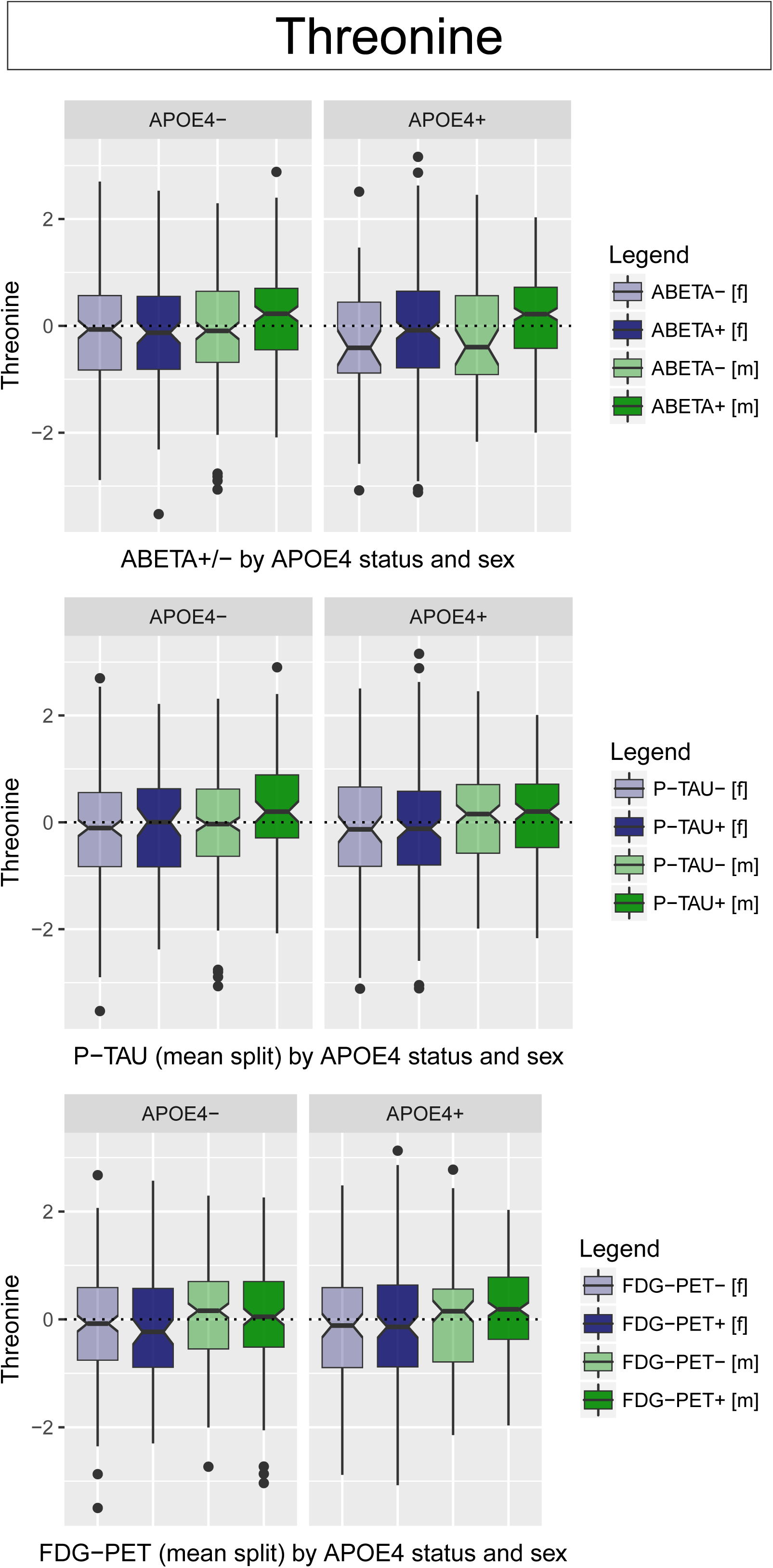
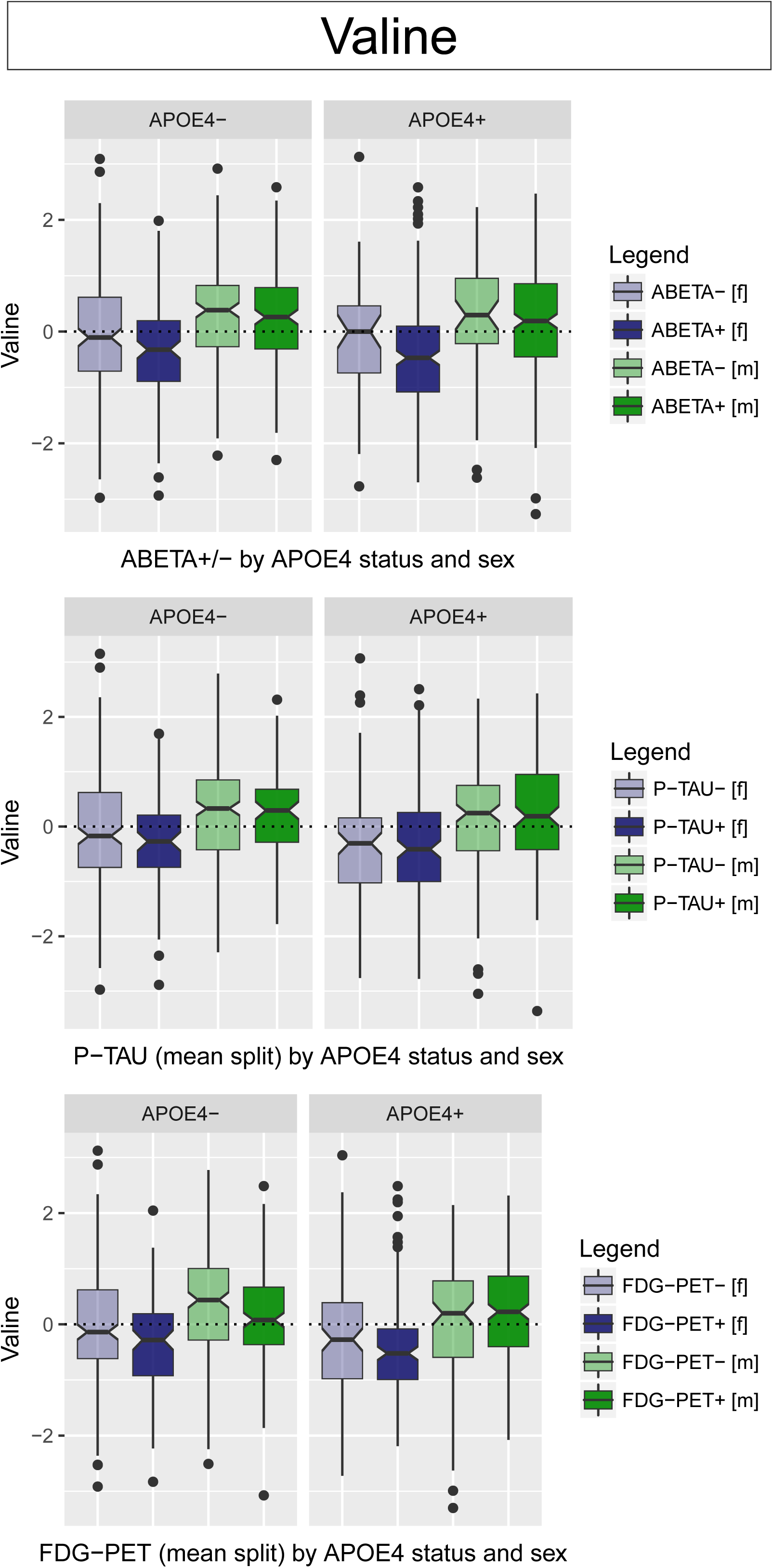
Boxplots for all 21 metabolites identified in this study in relation to A-T-N biomarkers in 2-fold stratified analyses In a separate file, we provide boxplots for all 21 metabolites identified in this study to show their relation to A-T-N biomarkers (A: pathological CSF Aβ_1-42_; T: mean-split CSF p-tau levels; N: mean-split FDG-PET values) for 2-fold stratified analyses by both sex and *APOE* ε4 status. *APOE* ε4 status groups are plotted in separate panels, females and males are distinguished by color (f: blue, m: green), and binarized biomarker groups are emphasized by lighter (lower-risk biomarker profile) and deeper (higher-risk biomarker profile) colors.

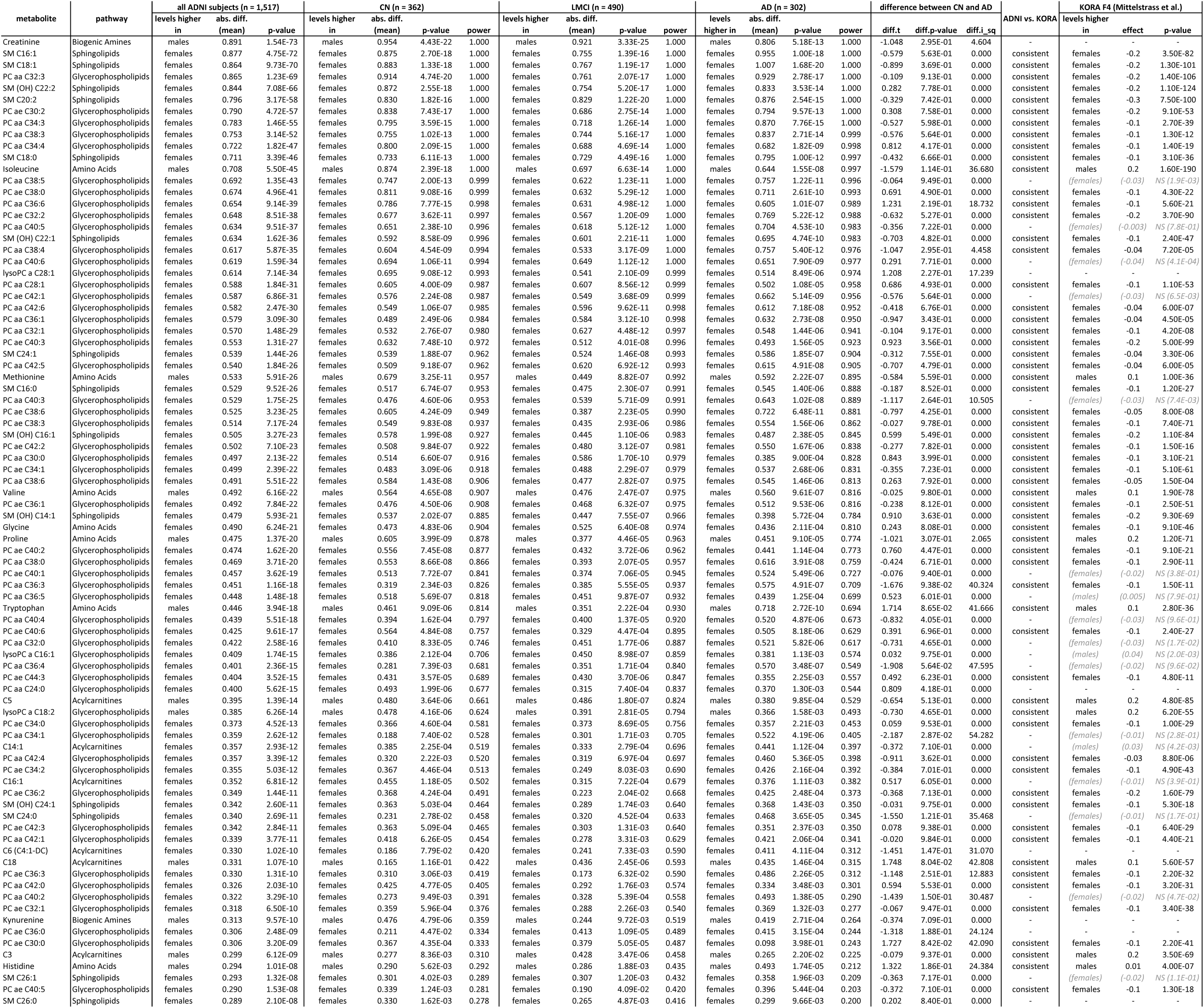
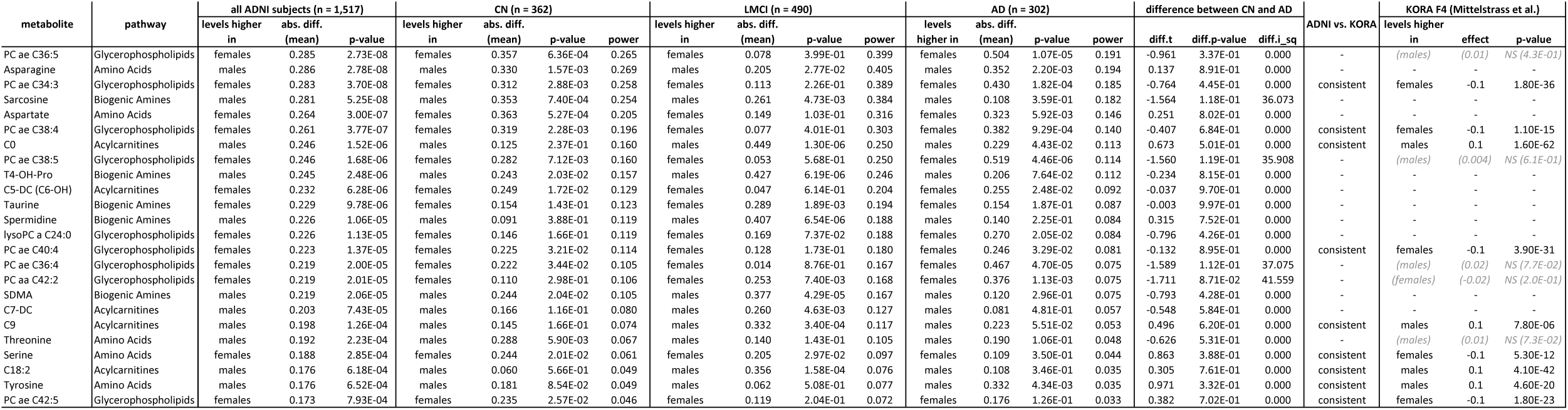
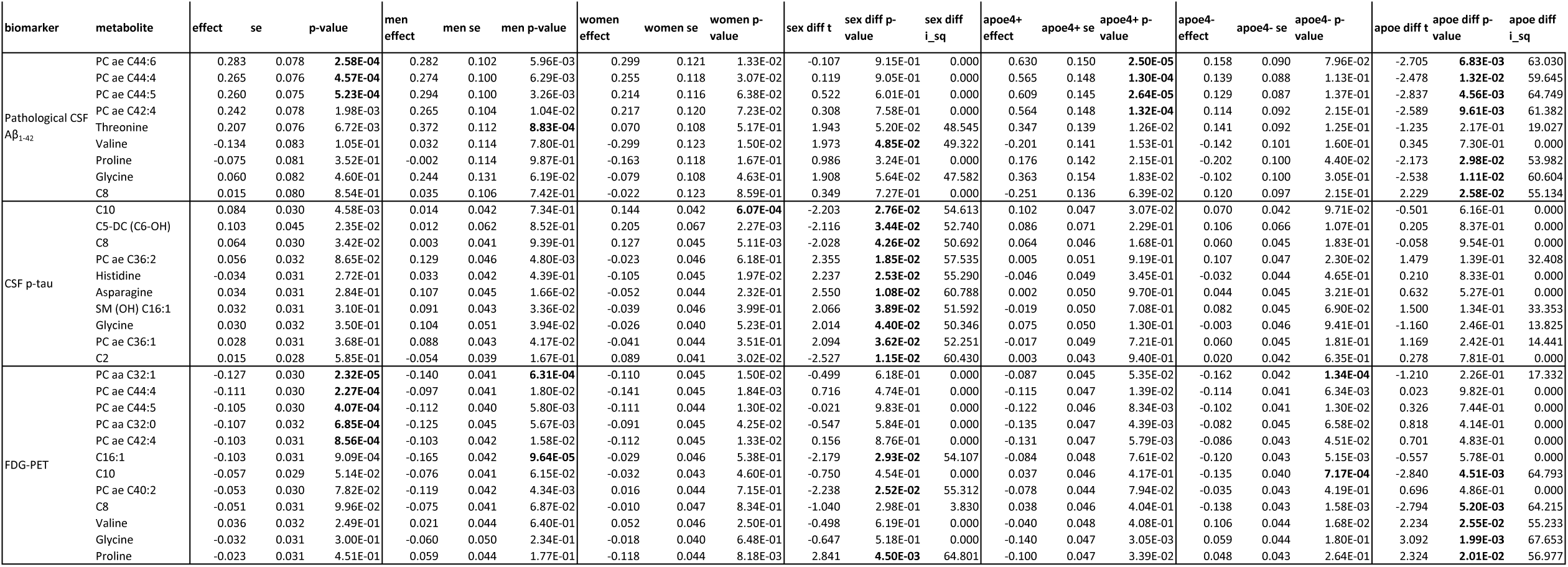

