## Supplemental Text for "The Alzheimer’s Disease Metabolome: Effects of Sex and *APOE* ε4 genotype"

### Data Availability

Metabolomics datasets from the Biocrates p180 platform used in the currently analyses for the ADNI-1, AGNI GO/2, and ROSMAP cohorts are available via the Accelerating Medicines Partnership-Alzheimer’s Disease (AMP-AD) Knowledge Portal and can be accessed at <http://dx.doi.org/10.7303/syn5592519> (ADNI-1), <http://dx.doi.org/10.7303/syn9705278> (ADNI GO/2), and <http://dx.doi.org/10.7303/syn10235592> (ROSMAP). The full complement of clinical and demographic data for the ADNI cohorts are hosted on the LONI data sharing platform and can be requested at [http://adni.loni.usc.edu/data-samples/access-data/](https://urldefense.proofpoint.com/v2/url?u=http-3A__adni.loni.usc.edu_data-2Dsamples_access-2Ddata_&d=DwMGaQ&c=imBPVzF25OnBgGmVOlcsiEgHoG1i6YHLR0Sj_gZ4adc&r=nsCVdiiwgH4TuvJab43dDXBo913IGTsJ7EODyQSlpaLT_T7XkmGJDiPo7F4n_q73&m=NeeaDvp_MwOm_WUkFLF9HdtygDg5xatnn2U7adVW_dE&s=ioYyc_1eCamqWj70WRGXnBgOfxij-8L_eZ0jrIsAWQw&e=)  .  The full complement of clinical and demographic data for the ROSMAP cohorts are available via the RUSH AD Center Resource hub and can be requested at [https://www.radc.rush.edu/](https://urldefense.proofpoint.com/v2/url?u=https-3A__www.radc.rush.edu_&d=DwMGaQ&c=imBPVzF25OnBgGmVOlcsiEgHoG1i6YHLR0Sj_gZ4adc&r=nsCVdiiwgH4TuvJab43dDXBo913IGTsJ7EODyQSlpaLT_T7XkmGJDiPo7F4n_q73&m=NeeaDvp_MwOm_WUkFLF9HdtygDg5xatnn2U7adVW_dE&s=AN56JZOWJSBLUG2tRWB2I8yo_T95bIjpyjUN83fd_YM&e=) .
